# Kv1.2 contributes to pattern separation by regulating the hippocampal CA3 neuronal ensemble size

**DOI:** 10.1101/2020.07.21.213538

**Authors:** Kisang Eom, Hyoung Ro Lee, Jung Ho Hyun, Hyun-Hee Ryu, Yong-Seok Lee, Inah Lee, Won-Kyung Ho, Suk-Ho Lee

**Affiliations:** Cell Physiology Lab. Department of Physiology, Seoul National University College of Medicine; Department of Brain and Cognitive Science, Seoul National University College of Natural Science, 103 Daehak-ro, Jongno-gu, 03080, Seoul, Republic of Korea

## Abstract

Kv1.2 expression in rodent CA3 pyramidal cells (CA3-PC) is polarized to distal apical dendrites, and regulate the synaptic responses to perforant pathway (PP) inputs. Accordingly, Kv1.2 haploinsufficiency (*Kcna2*+/−) in CA3-PCs, but not Kv1.1 (*Kcna1*+/−), lowered the threshold for long-term potentiation at PP-CA3 synapses. The *Kcna2*+/− mice, but not *Kcna1*+/−, displayed impairments in contextual fear discrimination task. The size and overlap of CA3 ensembles activated by the first visits to slightly different contexts were not different between wildtype and *Kcna2*+/− mice, but these ensemble parameters diverged over training days between genotypes, suggesting abnormal plastic changes in the CA3 network of *Kcna2*+/− mice. Eventually, the *Kcna2*+/− mice exhibited larger ensemble size and overlap upon retrieval of two contexts, compared to wildtype or *Kcna1*+/− mice. These results suggest that Kv1.2 subunits prevent promiscuous plastic changes at PP-CA3 synapses, and contribute to sparse representation of memories and pattern separation in the CA3 network.

## Introduction

Hippocampal CA3 region is implicated in rapid formation of episodic memories (Kesner, 2013). Synaptic inputs from entorhinal cortex (EC) converge on hippocampal CA3 pyramidal cells (CA3-PCs) directly via perforant pathway (PP) and indirectly through dentate gyrus (DG) via mossy fibers (MFs). DG is thought to segregate overlapping input patterns (pattern separation) by expansion re-coding of input patterns on the large number of dentate granule cells (GCs) that exhibit sparse firings (Senzai and Buzsaki, 2016; GoodSmith et al., 2017). Because MFs sparsely innervate CA3-PCs (Amaral et al, 1990) and convey ensemble patterns pre-processed by DG to CA3, computational models propose that MF inputs contribute to discrete and decorrelated representation of memories in the CA3 network, indicative of pattern separation (Leutgeb et al., 2004; Rolls and Kesner, 2006; Leutgeb et al., 2007). In contrast to MFs, PP densely innervates CA3-PCs and makes weak and Hebbian synapses (Amaral et al., 1990; McMahon and Barrionuevo, 2002). A formal model implies that cued memory retrieval of a pattern stored in recurrent network requires afferent inputs mediated by large number of associatively modifiable synapses, which correspond to PP-CA3 synapses (Treves and Rolls, 1992). Moreover, PP inputs are expected to be more informative for retrieval than MF inputs, because noisy part of a cue from EC can be amplified by an intervening layer, DG (O’Reilly and McClelland, 1994). To avoid interference between patterns stored in recurrent network, each pattern should be represented by a small number of principal neurons (sparse coding; Treves & Rolls, 1992). Given that associational/commissural (A/C) synaptic inputs are weakened under a high cholinergic level (Vogt and Regehr, 2001), the model assumes that MF inputs dominate the firing pattern of CA3-PCs during a learning phase. <u>Because unit synaptic inputs incoming via PP are not strong sufficient to elicit somatic action potentials (APs) in the postsynaptic CA3-PCs, there would be low probability for PP synaptic inputs alone to trigger long-term potentiation (LTP) at PP-CA3 synapses.</u> During learning phase, memory representation in a CA3 network is considered to be formed by Hebbian LTP at PP and A/C synapses in a subset of CA3-PCs chosen by decorrelated DG inputs (McMahon and Barrionuevo, 2002; Mishra et al., 2016). A previous lesion study, however, showed that MF inputs are dispensable for retrieval of acquired memories, whereas MF inputs are required for the memory acquisition (Lee and Kesner, 2004; but see Bernier et al., 2017), implying that <u>PP inputs initiate retrieval of memory representations stored in CA3 network</u>. In order for PP inputs to activate a decorrelated CA3 ensemble on retrieval, LTP at PP synapses should be induced in a small number of CA3-PCs (sparse encoding) during hippocampal learning. For sparse encoding at PP synapses, induction of LTP at PP synapses should be tightly regulated, and thus it would be of crucial importance to keep the LTP threshold (the minimal strength of synaptic inputs required for LTP induction) high at PP-CA3 synapses.

The CA3 region displays the highest expression of Kv1.2 transcripts in the rodent hippocampus (http://mouse.brain-map.org/gene/show/16263). Somatodendritic expression of Kv1.2 is polarized to distal apical dendrites, at which PP synaptic inputs arrive (Hyun et al. 2015). Consistent with the notion that D-type K^+^ current (I_K(D)_) is activated by low voltage depolarization near the AP threshold (Storm, 1988), distal dendritic Kv1.2 subunits regulate the threshold for dendritic Na^+^ channel-dependent amplification of PP-evoked EPSPs (PP-EPSPs) (Hyun et al., 2015). Because PP synapses exhibit Hebbian plasticity, we hypothesized that downregulation of Kv1.2 and consequent enhancement of EPSP-to-spike (E-S) coupling would have a profound effect on the propensity of LTP induction at PP synapses. Here, we show that insufficiency of Kv1.2 in CA3-PCs, whether it is caused by activity-dependent downregulation or heterozygous knock-out of *Kcna2* (*Kcna2*+/−), <u>lowers the LTP threshold at PP-CA3 synapses</u>. The non-specific lowering of LTP threshold at PP-CA3 synapses caused by genetic haploinsufficiency of *Kcna2* was well correlated with abnormal enlargement of CA3 ensembles during contextual fear discrimination task and impaired pattern separation of CA3 ensembles representing slightly different contexts.

## Results

### The threshold for PP-LTP is lowered by haploinsufficiency of Kcna2

Homosynaptic LTP of PP-EPSCs (referred to as PP-LTP) in the CA3-PCs of wildtype (WT) mice was induced under the current clamp mode by high frequency stimulation (HFS) of PP, which consists of 10 bursts repeated every 10 s with each burst being comprised of 20 pulses at 100 Hz (PP-HFS, Figure 1Aa and Ab) (McMahon et al, 2002). We monitored PP-EPSCs of non-failure events before and after PP-HFS. <u>When PP-EPSC amplitudes were increased by more than 50% at 30 min after PP-HFS, it was regarded as induction of PP-LTP.</u> To confirm stimulation of PP, effects of DCG-IV, a group II mGluR agonist, on synaptic responses were tested at the end of experiments (Figure 1Ac) (Tsukamoto et al., 2003). The PP stimulation strength was quantified as the baseline amplitudes of PP-EPSC. <u>The PP-LTP threshold was determined by the minimal amplitude of baseline PP-EPSCs that induces PP-LTP</u>. At weak stimulation strength, which evoked the baseline EPSCs less than 12 pA, PP-HFS induced neither somatic firing nor PP-LTP (Figure 1Aa and open circles in Figure 1Ac; 8.48 ± 1.02 to 8.52 ± 0.95 pA, n = 5, t = −0.137, p = 0.898, paired t-test). We repeated the same experiments with stronger stimulation intensities, which elicited baseline EPSCs amplitudes larger than 12 pA. The strong PP-HFS always drove the postsynaptic CA3-PCs to fire APs (Figure 1Ab), and induced PP-LTP (black dots in Figure 1Ac; 14.61 ± 0.78 to 27.55 ± 2.72 pA, n = 5, t = −6.415, p = 0.003, paired t-test), indicative of the Hebbian property of PP synapses. The mean values for potentiation of EPSCs by the weak and strong PP-HFS are summarized in Figure 1Ad. After the CA3-PCs underwent PP-LTP, CA3-PCs responded to the same burst stimulation of PP (20 pulse at 100 Hz) with the significantly higher number of somatic spikes (Figure 1 – figure supplement 1), implying that PP-LTP may contribute to spike transfer from EC to CA3.

**Figure 1.**
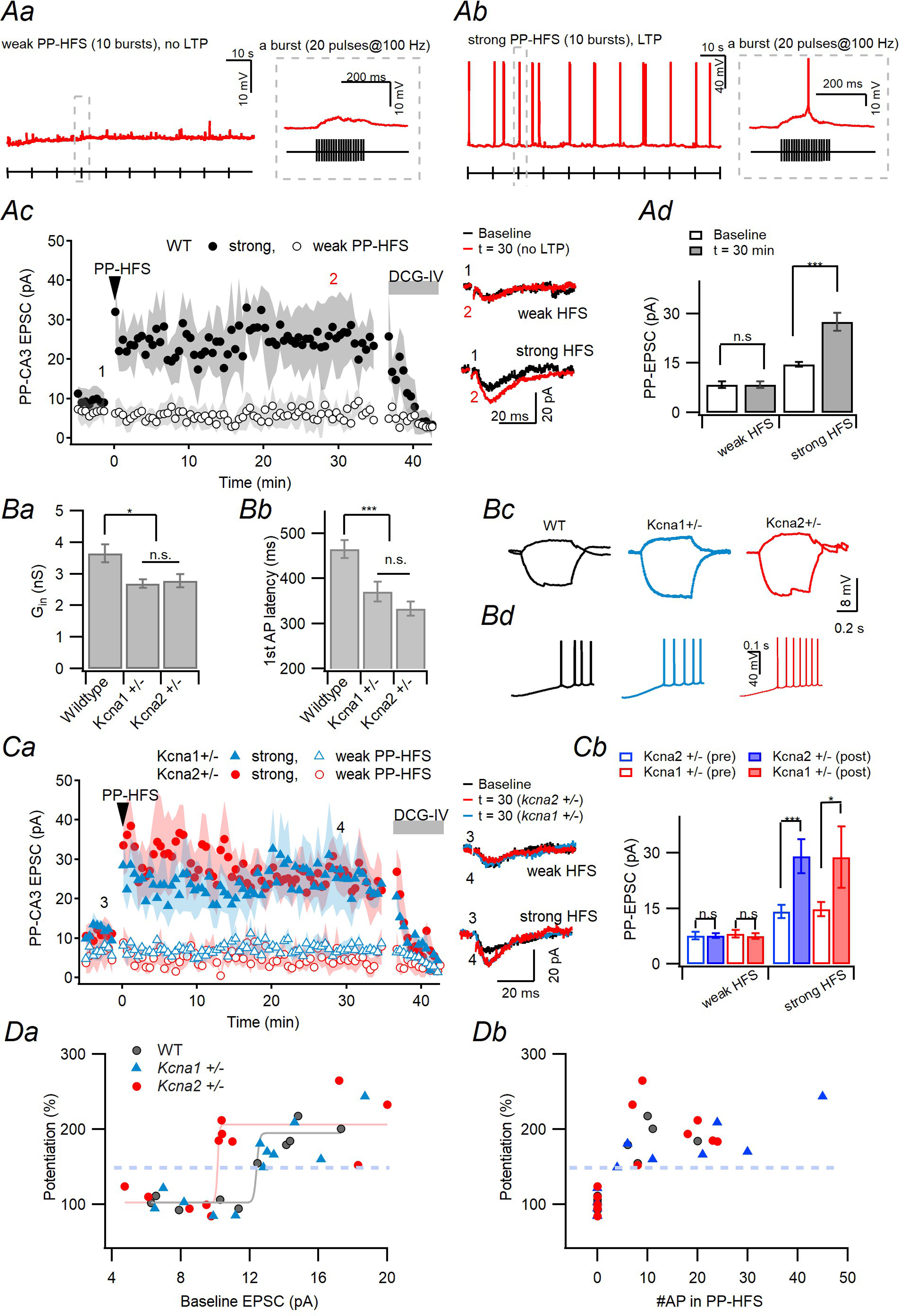
The threshold for homosynaptic LTP at PP-CA3 synapse (PP-LTP) is lower in *Kcna2+/−* than in WT and *Kcna1+/−* mice. **Aa.** Voltage responses of a CA3-PC (*red*) to weak HFS of PP (PP-HFS) protocol (*black*). PP-HFS is comprised of 10 bursts delivered every 10 s (*left*), and each bursts consists of 20 stimuli at 100 Hz (*right*, corresponds to the gray dashed box on the left). This HFS protocol was applied while the postsynaptic membrane potential was adjusted at -60 mV in current-clamp mode. **Ab.** Strong PP-HFS protocol. The pulse protocol was the same as the weak PP-HFS (*Aa*) but higher stimulation intensity. Note that a burst of APs was elicited by strong PP-HFS in the CA3-PC. **Ac**, Monitoring of PP-EPSCs before and after the PP-HFS in CA3-PCs of WT mice. Weak HFS and strong HFS were categorized based on the induction of PP-LTP. *Insets*: Representative PP-EPSC traces before (*black*) and 30 min after (*red*) weak (*upper*) or strong (*lower*) PP-HFS. **Ad.** Mean amplitude of EPSCs before and after weak or strong PP-HFS. All strong stimulation intensity elicited the baseline PP-EPSCs larger than 12 pA. **B**, Mean values for input conductance (G_in_, *Ba*) and first AP latency (*Bb*) measured in CA3-PCs of the three genotype mice. *Bc* and *Bd* show representative traces for voltage responses to subthreshold (+10/-30 pA) current injection (*Bc*), and those to ramp (250 pA/s for 1s) current injection (*Bd*), from which G_in_ and first AP latency were measured, respectively. Both parameters were lower in *Kcan1*+/− and *Kcna2*+/− than wildtype littermates (WT), but not different *Kcan1*+/− and *Kcna2*+/− CA3-PCs (G_in_, F_(2,35)_ = 7.539, p = 0.002; WT vs. *Kcna1+/−*, p = 0.003; WT vs. *Kcna2+/−*, p = 0.006; *Kcan1*+/− vs. *Kcna2*+/−, p = 1.00; For AP latency, F_(2,46)_ = 11.539, p = 0.002; WT vs. *Kcna1+/−*, p = 0.003; WT vs. *Kcna2+/−*, 0.006;, *Kcan1*+/− vs. *Kcna2*+/−, p = 1.00; one-way ANOVA and Bonferroni post-hoc test).. **Ca.** Monitoring of PP-EPSCs before and after weak (*open symbols*) or strong (*closed symbols*) PP-HFS in CA3-PCs of *Kcna1+/−* (*blue*) and *Kcna2+/−* (*red*) mice. The same color code was used in insets. *Insets*: Representative traces of PP-EPSCs before (black) and 30 min after weak (*upper*) and strong (*lower*) PP-HFS. **Cb.** Mean amplitude of EPSCs before and after PP-HFS. **Da**, PP-LTP levels as a function of stimulation intensity which was quantified as the baseline PP-EPSC amplitude. Note that the LTP threshold in *Kcna2+/−* (c.a. 10.4 pA) was lowered than that of *WT* or *Kcna1+/−* (c.a. 12.1 pA). **Db**, PP-LTP levels as a function of the number of somatic APs evoked by PP-HFS. Note that PP-LTP was always associated with somatic APs evoked by PP-HFS.

Because somatodendritic distribution of Kv1.2 is polarized to distal apical dendrites of CA3-PCs and downregulation of Kv1.2 in a CA3-PC enhances E-S coupling of PP synaptic inputs (Hyun et al., 2015), we tested whether haploinsufficiency of Kv1.2 has any effects on the threshold for induction of PP-LTP. Previously, it was confirmed that the D-type K^+^ current (*I*_K(D)_) in CA3-PCs of *Kcna2*+/− mice is reduced to c.a. 58% of that of their WT littermates (Hyun et al., 2015). *I*_K(D)_ is primarily mediated Kv1.1 and Kv1.2 subunits in central neurons (Guan et al, 2006), and both subunits are equipped with molecular domains required for axonal targeting (Gu et al., 2003; Gu et al., 2006). Previous studies indicate that Kv1.1 and Kv1.2 subunits are expressed in axonal and dendritic compartments of CA3-PCs, respectively (Sheng et al. 1994; Grosse et al. 2000; Hyun et al. 2015; Rama et al. 2017), while the soma of CA3-PCs express both subunits (Rama et al. 2017). Axonal expression of Kv1.2, however, is still obscure in CA3-PCs. Moreover, PP expresses both Kv1.1 and Kv1.2 (Monaghan et al., 2001). Therefore, we cannot rule out possible contributions of axonal Kv1.2 expressed in PP and/or in CA3-PCs to the PP-LTP threshold. To test this possibility, we compared the *Kcna1*+/− CA3-PCs with *Kcna2*+/− CA3-PCs for the PP-LTP threshold. The excitability of *Kcna1+/−* CA3-PCs was enhanced similar to *Kcna2*+/−. The intrinsic excitability was evaluated by two parameters: the input conductance (G_in_) and first spike latency (AP latency) (See Materials and Methods) (Hyun et al. 2013). Both of G_in_ and AP latency were lower in *Kcna1+/−* and *Kcna2+/−* CA3-PCs than their WT littermates, but not different between *Kcna1+/−* and *Kcna2+/−* (Figure 1B).

The LTP threshold and degree of potentiation were measured for *Kcna1*+/− and *Kcna2*+/− CA3-PCs using the same methods as in the WT mice (Figure 1C). Potentiation of PP-EPSCs estimated at 30 min after strong PP-HFS was not different between WT, *Kcna*1+/− and *Kcna2*+/− (Figure 1Ad and Cb; F_(2,16)_ = 0.881, p = 0.436, one-way ANOVA; WT, 187.0 ± 10.6%, n = 5; *Kcna1*+/−, 177.4 ± 16.8%, n = 5; *Kcna2*+/−, 203.1 ± 14.0%). The PP-LTP level was measured as a normalized amplitude of EPSCs 30 min after HFS relative to the baseline. The plot for the PP-LTP levels as a function of PP stimulation intensities, quantified as the baseline amplitude of PP-EPSCs, shows that the threshold for the induction of PP-LTP is left-shifted in *Kcna2*+/− (10.2 pA), compared to those of *Kcna1*+/− (12.8 pA) and WT (12.4 pA) (Figure 1Da), indicating that the expression level of Kv1.2 regulates the PP-LTP threshold. Whereas the LTP threshold was lower in *Kcna2*+/−, *Kcna2*+/− was not different from other genotypes in that PP-HFS always elicited somatic APs whenever it induced PP-LTP (Fig. 1Db).

### Activity-dependent downregulation of Kv1.2 lowers the threshold for PP-LTP

Insufficiency of Kv1.2 can be induced not only by genetic mutation but also by repetitive stimulation of afferent MFs or the soma of a CA3-PC. The latter results in a decrease in input conductance and enhanced E-S coupling of PP synaptic inputs, collectively referred to as long-term potentiation of intrinsic excitability (LTP-IE) (Hyun et al., 2013; Hyun et al., 2015; Eom et al., 2019). Notably, LTP-IE enhances PP-EPSPs but not PP-EPSCs, supporting the notion that it is attributed to intrinsic excitability changes of dendrites (Hyun et al., 2015). Given that *Kcna2* insufficiency lowers the threshold for PP-LTP, we tested whether LTP-IE induced by 10 Hz somatic stimulation also lowers the PP-LTP threshold (referred to as metaplasticity). Because *Kcna2*+/− as well as *Kcna2*−/− CA3-PCs lack LTP-IE (Hyun et al., 2015), and *Kcna2*−/− mice die within the 3^rd^ postnatal week (Brew et al., 2007), we examined the effects of LTP-IE on the PP-LTP threshold in WT, *Kcna1*+/− and *Kcna2+/−* CA3-PCs to confirm involvement of Kv1.2 in the metaplasticity. To this end, we adjusted the PP stimulation intensity which evokes EPSCs lower than 10 pA. At this intensity PP-HFS by itself does not induce PP-LTP in naïve CA3-PCs of the all three genotypes. Since PP-EPSPs are influenced by intrinsic excitability, EPSCs were monitored, and EPSPs were measured at the start and the end of EPSC monitoring. Somatic stimulation at 10 Hz did not alter PP-EPSCs (blue arrow in Figure 2Ac) in all three genotypes (Figure 2Ba), but it enhanced PP-EPSPs measured 6 min after priming in WT and *Kcna1+/−*, but not in *Kcna2+/−* (Figure 2Bb). To prevent washout of intracellular milieu necessary to induce LTP, the PP-HFS was delivered within 20 min after the patch break-in (Malinow and Tsien, 1990). Distinct from the naïve CA3-PCs (Figure 1Aa), subsequent weak PP-HFS (PP-EPSC < 10 pA) readily elicited AP firing in the WT and *Kcna1*+/− CA3-PCs which underwent 10 Hz somatic stimulation (Figure 2Aa), but such E-S potentiation did not occur in *Kcna2*+/− CA3-PCs (Figure 2Ab). Accordingly, weak PP-HFS induced PP-LTP in WT and *Kcna1*+/−, but not in *Kcna2*+/− CA3-PCs (Figure 2Ac) (F_(2,16)_ = 22.510, p < 0.001; WT vs. *Kcna1+/−*, p = 1.00; For WT or *Kcna1+/− vs*. *Kcna2+/−*, p < 0.001; one-way ANOVA, Bonferroni post-hoc test). Figure 2C summarizes the dependence of PP-LTP level on the baseline EPSC amplitudes. This plot demonstrates that 10 Hz somatic stimulation lowers the threshold for PP-LTP induction in WT and *Kcna1*+/− mice. In contrast, the threshold for PP-LTP in naïve *Kcna2+/−* CA3-PCs was already lower than other genotypes, and not affected by 10 Hz somatic stimulation. These results suggest that downregulation of Kv1.2 mediates both of LTP-IE and metaplasticity induced by 10 Hz somatic stimulation, but we cannot rule out a possibility that the 10 Hz somatic stimulation may lower the PP-LTP threshold by cellular mechanisms other than downregulation of Kv1.2. The effect of 10 Hz somatic APs on the LTP threshold was even larger than *Kcna2* haploinsufficiency, reflecting our previous observations that I_D_ in CA3-PCs was more reduced by 10 Hz somatic APs (to c.a. 30% of control; Hyun et al., 2013) than by *Kcna2*+/− (to c.a. 58%; Hyun et al., 2015).

**Figure 2.**
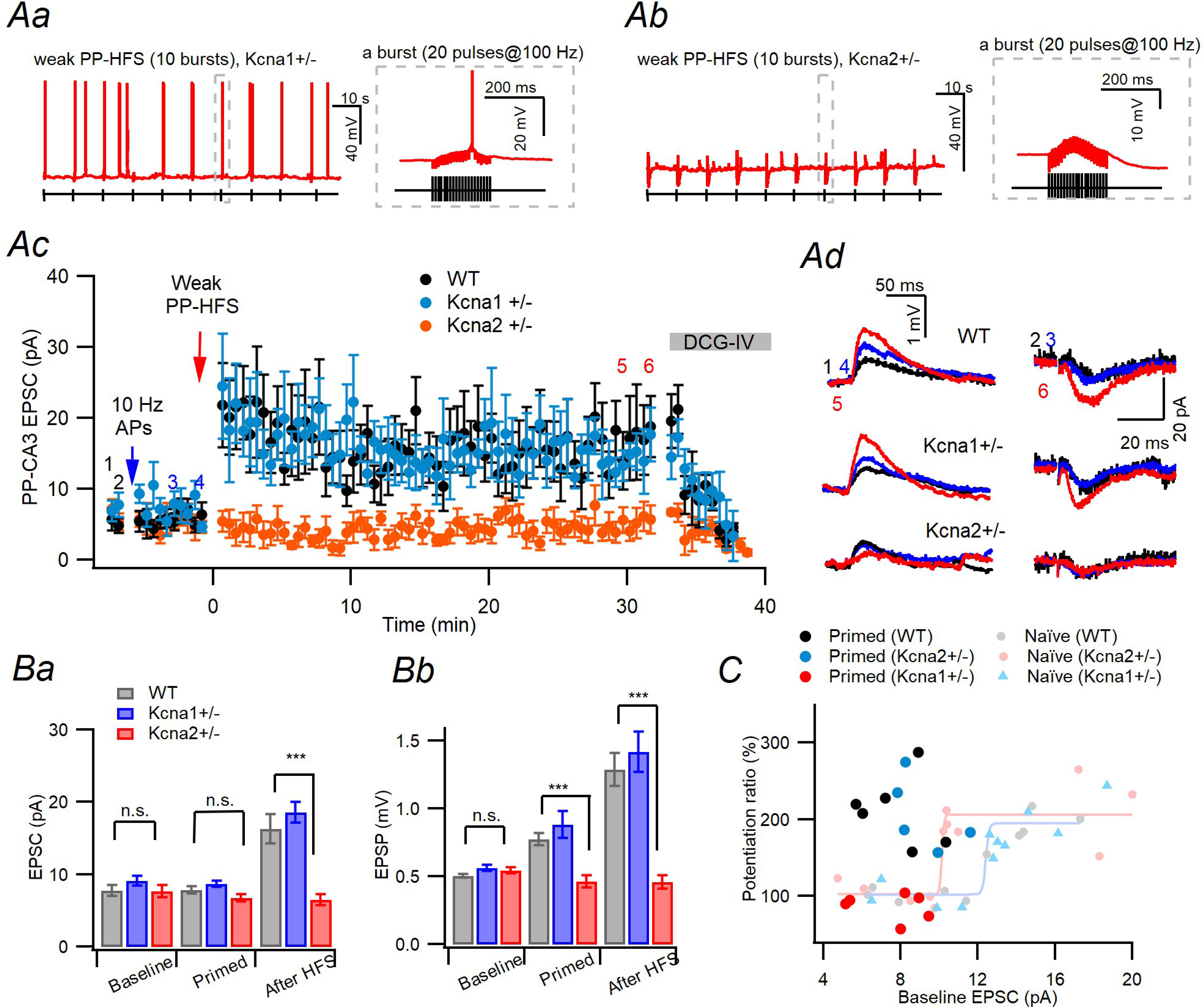
Kv1.2 mediates the 10 Hz somatic stimulation-induced metaplasticity at PP-CA3 synapses. **Aa-Ab**, Postsynaptic voltage responses to weak PP-HFS in *Kcna1*+/− (*Aa*) and *Kcna2*+/− (*Ab*) CA3-PCs that underwent 10 Hz somatic stimulation. Gray dashed box regions are shown in expanded time scale on the right insets. **Ac.** Metaplasticity induced by somatic conditioning at *WT* (*black*), *Kcna1+/−* (*blue*), and *Kcna2+/−* (*red*) PP-CA3 synapses. PP-EPSCs are monitored before and after somatic conditioning at -5 min (*blue arrow*) and subsequent weak PP-HFS at 0 min (*red arrow*). After the somatic stimulation, LTP was induced by weak HFS in WT and *Kcna1*+/−, but not in *Kcna2*+/− CA3-PCs. **Ad**. Representative traces for PP-EPSPs (*left*) and PP-EPSCs (*right*) recorded in wildtype (*upper*), *Knca1+/−* (*middle*), and *Kcna2+/−* (*lower*) CA3-PCs at three time points indicated in *Ac* by numbers (1,baseline, *black*; 4, after conditioning, *blue*; 5, after PP-HFS, *red*). **Ba-Bb**. Mean amplitudes for PP-EPSCs (*Ba*) and PP-EPSPs (*Bb*) recorded before (*Baseline*) and after (*Primed*) 10 Hz somatic stimulation, and 30 min after weak PP-HFS (*After HFS*) in the three genotype CA3-PCs. Note that somatic conditioning increased PP-EPSPs (WT, n =6, t = −6.086, p = 0.002; *Kcna1*+/−, n = 7, t = −3.642, p = 0.022; *Kcna2*+/−, n = 5, t = 1.851, p = 0.114; paired t-test) but not EPSCs (WT, t = −0.132, p = 0.900; *Kcna1*+/−, t = 0.518, p = 0.632; *Kcna2*+/−, t = 0.302, p = 0.773; paired t-test) in WT and *Kcna1+/−*. Neither EPSPs nor EPSCs was altered by conditioning in *Kcna2*+/−. Furthermore, PP-HFS failed the induction of PP-LTP. **C.** Dependence of the PP-LTP levels on the PP stimulation intensity quantified as the baseline PP-EPSC amplitudes in naïve (*light-colored*, adopted from Figure 1D) and primed CA3-PCs. Priming was induced by 10 Hz somatic stimulation. Note that no metaplasticity was induced in *Kcna2+/−* CA3-PCs by the priming.

It should be noted that 10 Hz somatic stimulation in Figure 2 was used as a surrogate of high frequency MF stimulation, and that MF inputs are privileged for the induction of LTP-IE (Eom et al., 2019). Indeed, we confirmed that repetitive stimulation of afferent MFs (20 Hz, 2 s) induces metaplasticity at PP synapses similar to 10 Hz somatic APs (Figure 2-figure supplement 1). Taken together, above results indicate that insufficiency of Kv1.2, whether it is induced by genetic mutation or by 10 Hz somatic APs, facilitates <u>induction of PP-LTP.</u> Assuming that the probability that PP inputs alone induce Hebbian plasticity is fairly low, our results together with McMahon et al (2002) imply that <u>induction of PP-LTP</u> may be allowed in the PCs receiving concurrent MF inputs and in PCs primed by previous strong MF inputs, both of which anyway depend on MF inputs.

### Kv1.2+/− mice, but not Kv1.1+/− mice, are impaired in rapid contextual discrimination

Given that PP-LTP enhances spike transfer from EC to CA3, lowered threshold for PP-LTP would potentially increase the size and overlap of CA3 ensembles. Both MF-induced LTP-IE and genetic *Kcna2* insufficiency have potential to exert adverse influence on pattern separation. In WT mice the PP-LTP threshold is lowered selectively in a CA3-PC primed by MF inputs, and thus only limited number of PCs would be primed owing to the sparseness of MF innervation and firings of dentate GCs. In contrast, PP-LTP threshold in *Kcna2*+/− mice is lowered not in a subset of CA3-PCs but over the whole CA3-PC population. Therefore, Kv1.2 insufficiency may increase the chance for MF input-independent induction of PP-LTP, and promiscuous encoding at PP synapses would ensue. We hypothesized that non-specific lowering of PP-LTP threshold caused by haploinsufficiency of Kv1.2 may result in impairment in a pattern-separation task, in which mice must learn association of different behaviors with two slightly different contexts.

Although *Kcna2−/−* mice had a reduced life span (3 weeks; Brew et al., 2007), we found that their heterozygous littermates (*Kcna2+/−*) had a normal life span. There were no changes in activity, feeding, reproductive, or parental behaviors in *Kcna1+/−* (n = 10) and *Kcna2+/−* (n = 9) mice, compared to WT (n = 11) mice. To examine behavioral pattern separation, the three genotype mice (WT littermates, *Kcna1+/−* and *Kcna2+/−*) were subject to the contextual fear discrimination task (McHugh et al., 2007; Kim et al., 2020). This task is comprised of three phases: contextual acquisition phase (day 1-3), generalization test phase (day 4-5), and the discrimination training phase (day 6-14) (Figure 3A). In the first phase, all genotype mice were given by a foot shock 180 s after being placed in context A (conditioned context) for consecutive 3 days. In the 2nd phase (day 4 and 5), the freezing behavior of such conditioned mice was assessed without a foot shock in context A and context B (safe context). The context B had the same sound noise level, metal grid floor, sidewalls and roof as in context A, but differed from context A in having a unique odor (1% acetic acid), dimmer light (50%) and a slanted grid floor by 15° angle (Figure 3-figure supplement 1). On the subsequent 9 days (day 6 -14, phase 3), mice daily visited both contexts with a foot shock only in context A (Figure 3A). The kinetics of freezing in context A was not different among the three genotypes (Figure 3Ba, Table 1 for statistics). All genotypes showed similar levels of freezing behavior in context B as in contexts A suggesting that all genotypes displayed equivalent generalization on day 4 and 5 (Figure 3Bb, Table 1). WT and *Kcna1*+/− mice began to distinguish the two contexts on the 2nd day of phase 3 (day 7), as shown by the increase in the discrimination ratio [freezing in A/(A+B)] (Figure 3Bc). *Kcna2+/−* mice, however, exhibited significant delay in the acquisition of the discrimination (Figure 3Bc, Table 1; WT *vs*. *Kcna1+/−*, p = 1.00; For WT or *Kcna1+/− vs*. *Kcna2+/−*, p < 0.001). On day 9, the *Kcna2+/−* mice exhibited elevated freezing level not only in A but also in B, whereas WT and *Kcna1+/−* mice showed significantly lower freezing in context B (Figure 3Bd, Table 1; WT vs. *Kcna1+/−*, p = 0.135; For WT or *Kcna1+/− vs*. *Kcna2+/−*, p < 0.001; 2-way ANOVA, Bonferroni post-hoc test). On the last day of the discrimination training phase (day 14), all mice discriminated the two contexts to a similar degree (Figure 3Be), suggesting that normal expression of Kv1.2 channel subunit is required for rapid pattern separation between slightly different contexts.

**Figure 3.**
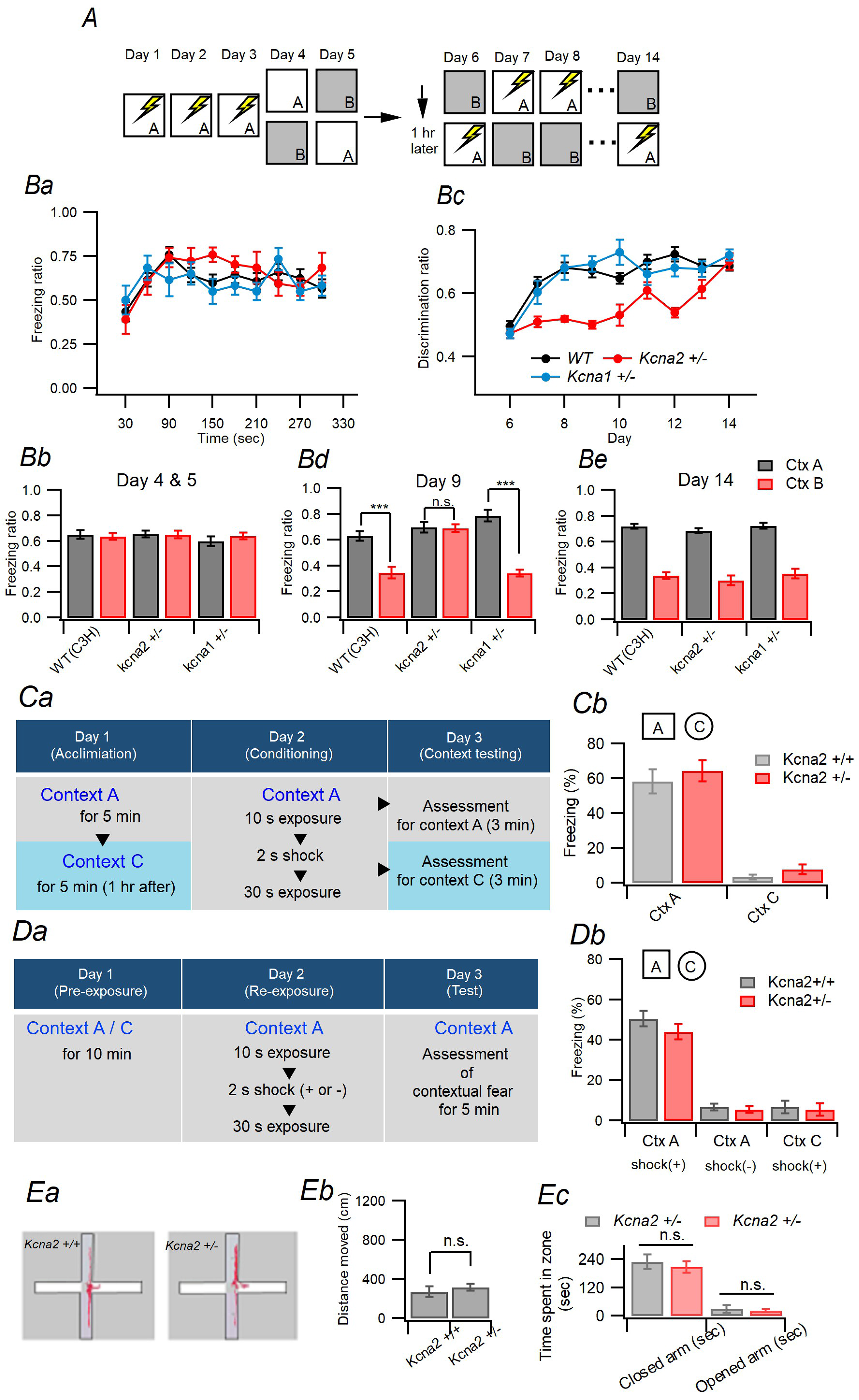
Kv1.2 is essential for rapid contextual discrimination. **A.** Protocol for contextual fear discrimination task. Each of the first 3 days, mice visited context A with receiving a single footshock (2 s, 0.75 mA). Over the subsequent 2 days (Day 4-5), freezing was measured in chamber A and B. During days 6 to 14, mice daily visited the two context with an hour interval with receiving a shock in A. Freezing behavior was assessed during the first 3 min in each context. **Ba.** Kinetics of freezing behavior over 5 min test in context A on day 4 and 5. **Bb.** Freezing ratio on day 4 and 5 of WT, *Kcna1+/−* and *Kcna2+/−* mice in context A (*gray*) and B (*red bars*). All genotype mice could not discriminate two contexts. **Bc**. Daily improvements in discrimination ratio over the phase 3 (day 6 -14) estimated in the three genotype mice. **Bd-Be.** Freezing ratio on day 9 (*Bd*) and 14 (*Be*) of three genotype mice in context A (*gray*) and B (*red bars*). On day 9, comparing the freezing ratio for A vs. B context in each genotype, WT: p < 0.001, *Kcna1*+/−: p < 0.001, *Kcna2*+/−: p = 0.914 (two-way ANOVA and simple effect analysis). **Ca.** Protocol for one-trial contextual fear conditioning, in which the test context C is very distinct from the conditioning context A. **Cb.** Freezing ratio of *Kcna2*+/− and their littermate *Kcna2*+/+ mice in context A and C measured 24 hours after conditioning in context A with a single 0.75 mA foot shock for 2 s (genotype, F_(1,19)_ = 0.137, p = 0.716; Context, F_(1,19)_ = 133.125, p < 0.001; genotype × context, F_(1, 19)_ = 0.007, p = 0.933, two-way ANOVA). **Da.** Protocol for pre-exposure mediated contextual fear conditioning (PECFC). **Db.** Freezing levels for context A and C measured 1 day after re-exposure for the context A (genotype, F_(1,46)_ = 0.286, p = 0.596; context, F_(1,46)_ = 77.173, p < 0.001; shock, F_(1,46)_ = 77.173, p < 0.001, 3-way ANOVA). **Ea.** Exemplar traces of WT and *Kcna2*+/− mice in the elevated plus maze (EPM) for a 5 min session. **Eb.** Total distance moved in the EPM for WT (n=5) and *Kcna2*+/− (n=10) mice was not different significantly (t = 0.691, p = 0.502, independent t-test). **Ec.** The time spent in closed arms (t = − 0.539, p = 0.599) and opened arms (t = −0.421, p = 0.680, independent t-test) for each animal were not different significantly.

**Table 1.**
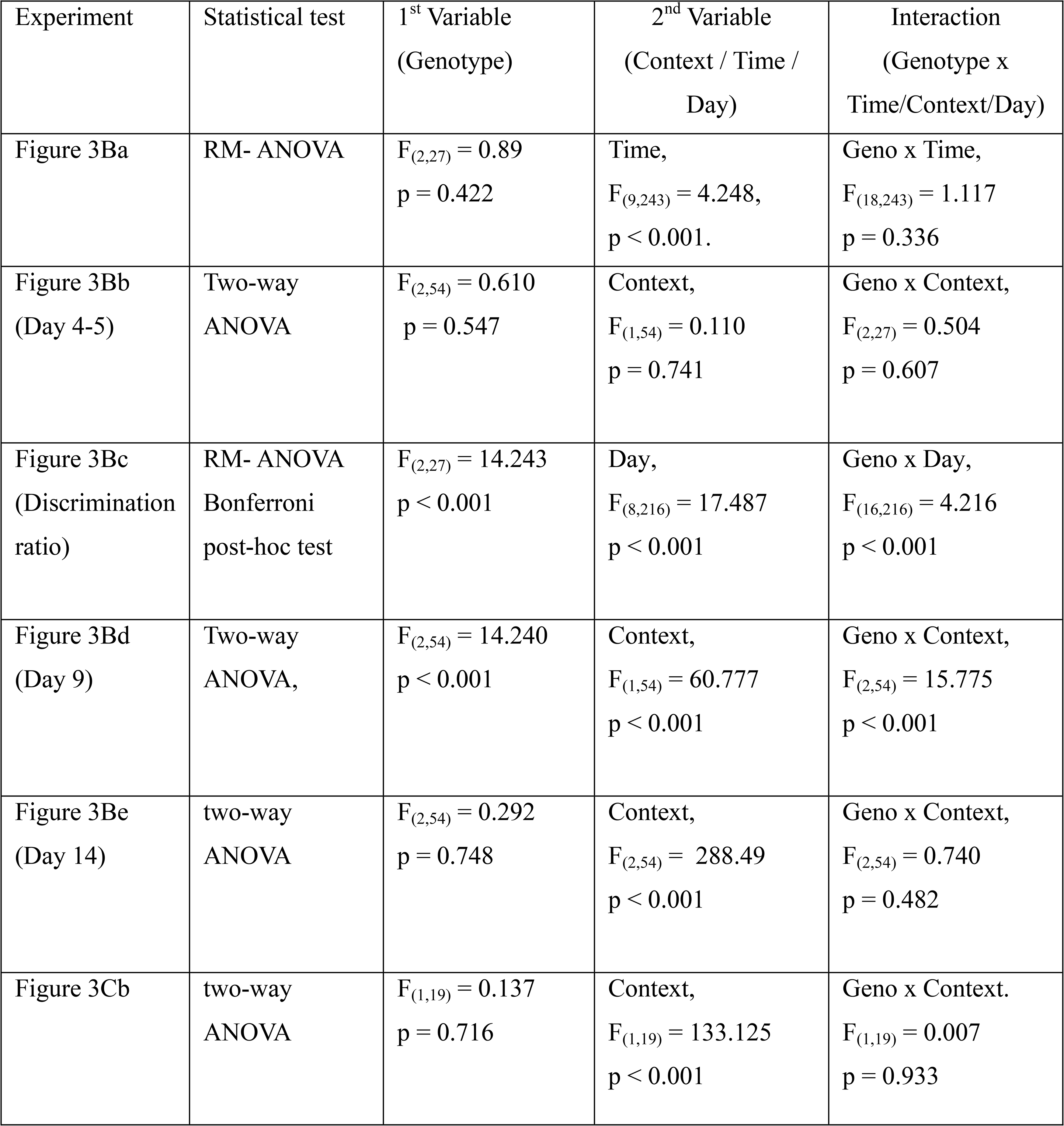
Statistical test for experiments shown in Figure 3

To rule out possible impairment in basic hippocampal contextual learning in *Kcna2+/−* mice, WT and *Kcna2+/−* mice were assessed for fear acquisition and retrieval of contextual freezing behavior based on Cravens et al. (2006). We examined the context specificity for fear conditioning by assessment of freezing behavior in two distinct contexts (A and C) 24 hours after pairing context A with a shock (Figure 3Ca). For context A, both genotype mice acquired and retained the fear conditioning to a similar degree after a single context-footshock pairing. The distinct context (context C), however, caused significantly lower freezing levels than the conditioned context in both genotype mice (Figure 3Cb, Table 1).

Next, we examined the capability of WT and *Kcna2+/−* mice for pattern completion-based memory retrieval using the pre-exposure-mediated contextual fear conditioning (PECFC) paradigm (Fanselow, 1990; Nakashiba et al., 2008) (Figure 3Da). The PECFC task requires retrieval of contextual memories from a very brief exposure to previously experienced context, which is impaired by disruption of synaptic output from CA3-PCs (Nakashiba et al. 2008). We found no difference between WT and *Kcna2*+/− mice in freezing response under the PECFC paradigm (Figure 3Db) (genotype, F_(1,46)_ = 0.286, p = 0.596; context, F_(1,46)_ = 77.173, p < 0.001; shock, F_(1,46)_ = 77.173, p < 0.001, 3-way ANOVA), implying that the *Kcna2+/−* mice are normal in CA3-dependent pattern completion-based retrieval. Finally, we assessed basal anxiety level using an elevated plus maze (EPM). Exemplar trails of WT, *Kcna2+/−* mice on the EPM (tracking for 5 min; Figure 3Ea). The total distances that mice explored in EPM was not significantly different between WT and *Kcna2+/−* mice (Figure 3Ea and 3Eb). The mean values for time spent in the closed and open arms were not significantly different between two genotype mice (Figure 3Ec).

### Kv1.2, but not Kv1.1, is crucial for pattern separation in the CA3

Above results indicate that *Kcna2*+/− mice are impaired in rapid discrimination of the contexts A and B tested in Figure 3A. We hypothesized that the impaired contextual discrimination in *Kcna2+/−* mice may be caused by larger overlap between neuronal ensembles representing the two slightly different contexts in the hippocampal CA3 region. To test this hypothesis, we examined CA3 cell ensembles activated upon memory retrieval of contexts A and B for the three genotype mice. An ensemble of active cells in each context was detected using catFISH (cellular analysis of temporal activity by fluorescence *in situ* hybridization) of immediate early genes (IEG) transcripts, Arc and Homer1a (H1a), which are expressed after patterned neuronal activities associated with synaptic plasticity (Guzowski et al., 2005). Designing Arc/H1a RNA probes targeting different locations from transcriptional initiation sites [22 bases for Arc and 51.6 kilobases (exon 5) for H1a] allows us to detect intra-nuclear foci (INF) of Arc/H1a transcripts at different timing from cellular activities (Vazdarjanova et al., 2002; Vazdarjanova and Guzowski, 2004). Typically, INF of Arc and H1a transcripts are expressed 5 and 30 min after cellular activity, respectively (Vazdarjanova et al., 2002). We confirmed the previous notion that most CA3-PCs are devoid of INF of H1a and Arc transcripts detected by catFISH 5 min or 30 min after an animal was exposed to a novel context, respectively (Figure 4 - Supplement 1).

As a control, we estimated an overlap between CA3 ensembles activated upon acquisition and retrieval of context A. To this end, WT and *Kcna2*+/− mice were handled daily for a week before the experiment to habituate to the general handling procedure. Little expression of c-fos was confirmed in the CA3 region of the handled mice compared to the mice that visited context A (Figure 4 - Supplement 2). The handled WT and *Kcna2*+/− mice visited the context A twice for 4 min with a 20 min interval, during which mice were kept in their home cage (Figure 4Aa). In the first visit, the mice received a footshock 3 min after being placed. After the second visit, mice were sacrificed, and their hippocampi underwent the Arc/H1a catFISH procedure. In the exemplar catFISH images of WT and *Kcna2*+/− CA3 cells (Figure 4Ab-Ac), H1a (green) and Arc (red) INF represent transcripts expressed upon 1^st^ and 2^nd^ visits, respectively. The fractions of H1a (+) and Arc (+) CA3 cells were not significantly different between WT and *Kcna2*+/− mice, suggesting that the increased distal dendritic excitability in *Kcna2*+/− CA3-PCs has little effect on the ensemble size activated by a novel context (For WT, H1a: 7.34 ± 0.49%, Arc: 7.21 ± 0.37%, n = 12 slices from 4 mice; For *Kcna2*+/−, H1a: 6.66 ± 0.83%, Arc: 6.61 ± 0.35%, n = 6 slices from 3 mice; 589 ± 92 cells per a slice; genotype, F_(1,36)_ = 0.419, p = 0.522; context, F_(1,36)_ = 2.797, p = 0.104; genotype × context, F_(1, 36)_ = 0.154, p = 0.697, two-way ANOVA; Figure 4Aa). The overlap between H1a (+) and Arc (+) ensembles was 3.70 ± 0.32% for WT and 3.00 ± 0.34% for *Kcna2*+/− (cells indicated by yellow arrowhead in Figure 4Aa; t = 1.363, p = 0.192, independent t-test). The conditional probability for re-activation of ensemble cells of the 1^st^ visit (A_1_) among ensemble cells of the 2^nd^ visit (A_2_) [P(A_1_|A_2_)] was not different between the two genotypes either (WT, 51.21 ± 3.47%; *Kcna2*+/−, 45.9 ± 3.11%).

**Figure 4.**
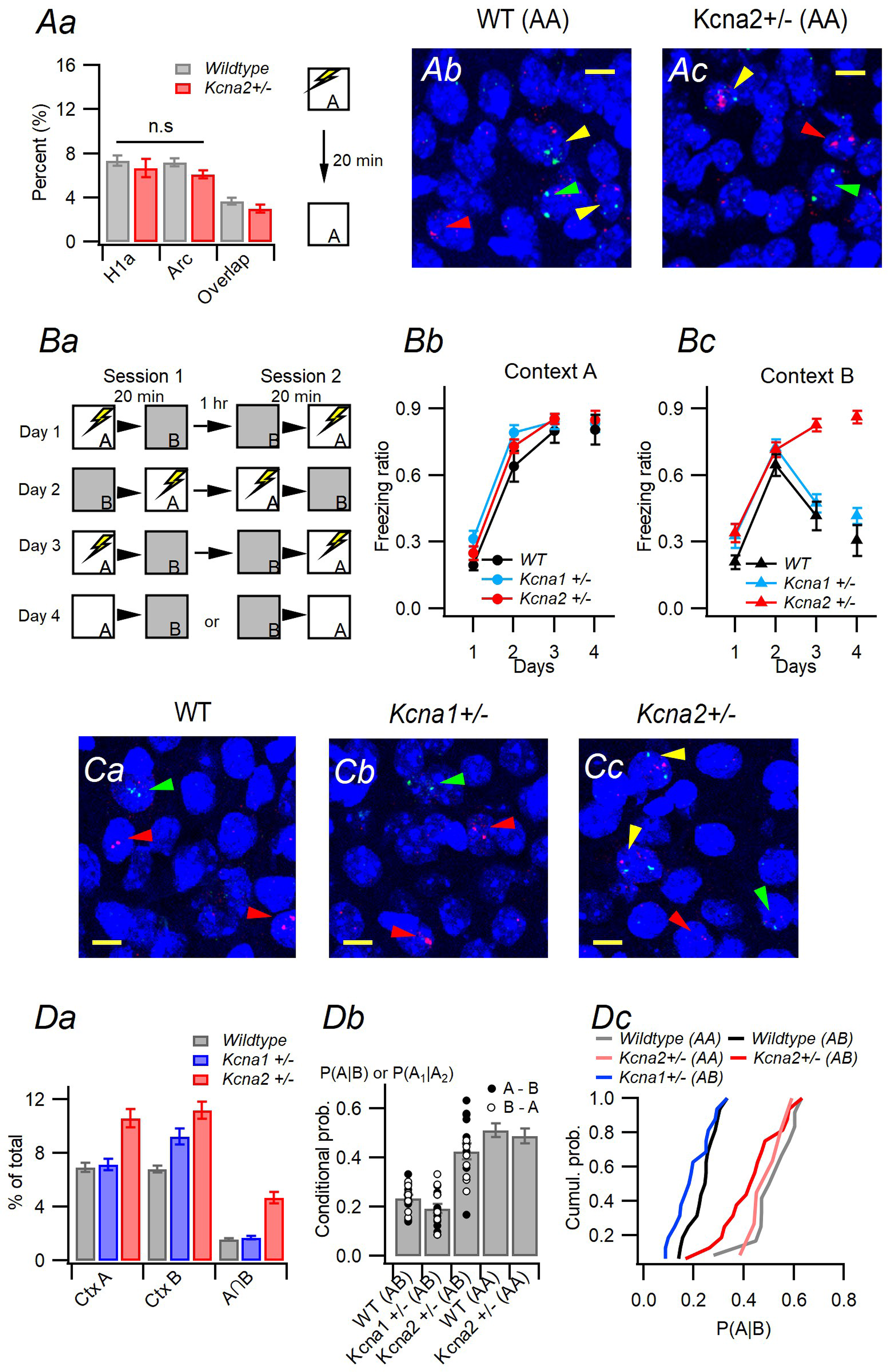
Insufficiency of Kv1.2 leads to impaired pattern separation of CA3 neuronal ensembles representing two similar contexts. **Aa.** Proportion of H1a(+), Arc(+) and H1a/Arc(+) (*overlap*) cells in the CA3 from *WT* (gray) and *Kcna2*+/− (red) mice, which explored the same context A twice with an interval of 20 min. On the 1^st^ visit, mice received a footshock. **Ab-c.** An exemplar fluorescence image of H1a and Arc transcripts in CA3 cells in *WT* (*Ab*) and *Kcna2*+/− (*Ac*) mice. Scale bars,10 μm. **Ba.** Protocol for modified contextual fear discrimination (CFD) test. For first 3 days, mice daily visited two similar contexts with 20 min interval with receiving a footshock always in context A. Mice were assessed for freezing for first 3 min after being placed in each context. On day 4, mice were allowed to freely explore the context A or B for 4 min (epoch 1). After a 20 min rest period in the home cage, mice were exposed to other context for 4 min (epoch 2). In 5 min after epoch 2, mice were killed for catFISH. **Bb-Bc.** Freezing ratio of mice in context A (Bb) and B (Bc) over 4 days, during which mice underwent the modified CFD test. WT and *Kcna1+/−* mice begun to discriminate two contexts in day 3, but *Kcna2+/−* mice could not even on day 4. **C.** Representative confocal images of *H1a* and *Arc* transcripts in nuclei (*blue,* counterstained with DAPI) of the CA3 pyramidal layer from WT (*Ca*), *Kcna1+/−* (*Cb*), and *Kcna2+/−* (*Cc*) mice. Nuclei expressing H1a alone, Arc alone, and both are indicated by green, red, and yellow arrowheads, respectively. *H1a* (*green dots*) and *Arc* (*red*) intra-nuclear foci are activated during epoch 1 and 2, respectively. Scale bars,10 μm. **Da.** Size of neuronal ensembles in CA3 activated in context A and B quantified as percentage of H1a(+) or Arc(+) cells among total cells in the CA3 pyramidal layer of each slice. The fraction of CA3 cells active in both of contexts A and B (denoted as A∩B) was not different between WT and *Kcna1*+/− mice, but higher in *Kcna2+/−* than other genotypes. The size of neuronal ensembles in *Kcna2+/−* was larger than that in *WT* and *Kcna1+/−.* **Db.** The conditional probability for ensemble cells active in context A among those active in context B [denoted as P(A|B)]. For comparison, conditional probability for re-activation of WT ensemble cells of the 1^st^ visit of context A (A_1_) among ensemble cells of the 2^nd^ visit (A_2_) [P(A_1_|A_2_)] is shown as WT(AA) and *Kcna2*+/−(AA). **Dc.** Cumulative probability histogram of the P(A|B) values for different genotypes. The overlap index between two ensembles in *Kcna2+/−* mice was significantly greater than that in *Kcna1+/−* or WT mice.

Next, we estimated the proportion of ensemble cells re-activated upon retrieval of contexts A and B in the same mice. After handling for a week, mice were subject to a contextual fear discrimination (CFD) protocol as shown in Figure 4Ba. Mice daily visited twice each of contexts A and B with receiving a footshock always in context A and never in B. The configurations of context A and B were the same as in Figure 3A (see Materials and Methods). On day 3, WT (n = 4) and *Kcna1+/−* (n = 4) mice began to distinguish the two contexts, whereas *Kcna2+/−* (n = 4) mice did not as shown by the difference in the freezing ratio (Figure 4Bb-Bc; Table 2 for statistics). On day 4, each genotype mice were divided into two subgroups. One subgroup was exposed to context A for 4 min with no footshock, returned to its home cage for 20 min, and then exposed to context B for 4 min (A/B). The other subgroup was subject to the same protocol except switching the order of A and B (B/A). On day 4, *Kcna2+/−* mice still did not distinguish the two contexts, in contrast to WT and *Kcna1+/−* (Figure 4Bb-Bc, Table 2). After visiting the second context, the mice were killed, and their hippocampi were examined for expression of Arc/H1a transcripts. Figure 4C shows exemplar Arc/H1a catFISH images of CA3 regions of the three genotype mice. For parameters of ensemble size and overlap, no difference was found between the two subgroups (A/B and B/A), and thus results are presented with respect to the context in Figure 4D. The fraction of CA3 cells activated in context A and that in context B were about 1.5 times larger in *Kcna2+/−* compared to WT (p < 0.001), and those in *Kcna1+/−* mice were marginally larger than WT (p = 0.053) (Table 2 and 3, Figure 4Da). The fraction of CA3 cells active in both of contexts A and B [denoted as P(A∩B)] was not different between WT and *Kcna1*+/− mice, but it was about three times higher in *Kcna2+/−* than other genotypes (Table 2 and 3). To test whether larger ensemble size in *Kcna2*+/− is responsible for larger overlap, we calculated the conditional probability for reactivation of the A ensemble cells among the B ensemble cells [P(A|B)] as an overlap index. The overlap index for *Kcna2+/−* was significantly higher than those of *Kcna1+/−* and WT (Table 2, Figure 4Db), and rather similar to the value for WT and *Kcna2*+/− mice which visited context A twice [denoted as WT(AA) and *Kcna2*+/−(AA), respectively] [WT(AA) vs. *Kcna2*+/−(AB), p = 0.188; *Kcna2*+/−(AA) vs. *Kcna2*+/−(AB), p = 1.00; one-way ANOVA, Bonferroni post-hoc test; Figure 4Db and Table 2]. The cumulative distribution of the overlap index shows a significant shift of the curve to the right in *Kcna2+/−*(AA), *Kcna2*+/−(AB) and WT(AA), compared to WT(AB) and *Kcna1+/−* [WT(AA) vs. WT(AB), Z = 2.4, p < 0.001; WT(AA) vs. *Kcna1+/−*, Z = 2.4, p < 0.001; WT(AA) vs. *Kcna2+/−*(AB), Z = 1.255, p = 0.083; WT(AA) vs. *Kcna2*+/−(AA), Z = 0.667, p = 0.766, Kolmogorov-Smirnov test, Figure 4Dc), indicating larger overlap between A and B ensembles in *Kcna2*+/−. If the overlap index of *Kv1.2+/−* were the same as that of WT [WT(AB), 23.6 ± 1.4%], the expected overlap between A and B ensembles should be 2.6% considering that P(B) in *Kcna2+/−* was 11.2 ± 0.7%. The measured value for overlap in *Kcna2+/−* was, however, 4.7 ± 0.4%, which is 1.8 times larger than the value expected under the same overlap index as WT. Therefore, larger overlap in *Kcna2+/−* cannot be explained by larger ensemble size alone, suggesting involvement of additional factors, probably less dependence of PP-LTP induction on concurrent MF inputs (see Discussion). Taken together with the results of Figure 3, pattern separation of CA3 ensembles are well correlated with the performance of CFD task across the three genotypes, implying that impaired pattern separation of CA3 ensembles may underlie the impairment of the rapid acquisition of contextual discrimination in *Kcna2*+/− mice.

**Table 2.** Statistical test for experiments shown in Figure 4

| Experiment | Statistical test | Genotype | Context | Genotype x Context | WT vs. *Kcna1*<sup>+/-</sup> | WT vs. *Kcna2*<sup>+/-</sup> | *Kcna2*<sup>+/-</sup> vs. *Kcna1*<sup>+/-</sup> |
| --- | --- | --- | --- | --- | --- | --- | --- |
| Figure 4Bb-Bc<br>Day 3 | two-way ANOVA, Bonferroni post-hoc test | $F_{(2,18)} = 20.817$<br>$p < 0.001$ | $F_{(1,18)} = 69.689$<br>$p < 0.001$ | $F_{(2,18)} = 13.91$<br>$p < 0.001$ | $p = 0.652$ | $p < 0.001$ | $p < 0.001$ |
| Figure 4Bb-Bc<br>Day 4 | | $F_{(2,18)} = 31.00$<br>$p < 0.001$ | $F_{(1,18)} = 86.22$<br>$p < 0.001$ | $F_{(2,18)} = 24.14$<br>$p < 0.001$ | $p = 0.292$ | $p < 0.001$ | $p < 0.001$ |
| Figure 4Da (Ensemble size) | | $F_{(2,96)} = 28.87$<br>$p < 0.001$ | $F_{(1,96)} = 0.27$<br>$p = 0.603$ | $F_{(2,96)} = 0.443$<br>$p = 0.643$ | $p = 0.053$ | $p < 0.001$ | $p < 0.001$ |
| Figure 4Da (Ensemble overlap) | one-way ANOVA | $F_{(2,48)} = 48.99$<br>$p < 0.001$ | irrelevant | irrelevant | $p = 1.00$ | $p < 0.001$ | $p < 0.001$ |
| Figure 4Db P(A\|B) | | $F_{(4,65)} = 33.50$<br>$p < 0.001$ | irrelevant | irrelevant | $p = 1.00$ | $p < 0.001$ | $p = 0.119$ |

**Table 3.** Mean values for size and overlap of neuronal ensembles activated by context A and B (mean ± SEM)

| Genotype (n=4 mice each) | P(A) (%) | P(B) (%) | P(A∩B) (%) | P(A)P(B) (%) | P(A\|B) (%) |
| --- | --- | --- | --- | --- | --- |
| WT | 7.00 ± 0.34 | 6.83 ± 0.27 | 1.56 ± 0.10 | 0.48 ± 0.04 | 23.2 ± 1.53 |
| *Kcna1*<sup>+/-</sup> | 7.15 ± 0.44 | 9.24 ± 0.61 | 1.69 ± 0.13 | 0.67 ± 0.06 | 19.3 ± 1.95 |
| *Kcna2*<sup>+/-</sup> | 10.6 ± 0.66 | 11.2 ± 0.65 | 4.69 ± 0.41 | 1.21 ± 0.12 | 42.6 ± 3.15 |

### CA3 ensembles of WT and Kcna2+/− mice differently evolve over training days

To further confirm the correlation between contextual discrimination and the overlap of CA3 ensembles, we examined the CA3 ensemble size and their overlap before WT and *Kcna2*+/− mice discriminate the contexts A and B. To this end, mice were subject to the same protocol of Figure 4Ba, but were killed for catFISH after one paired visits (session 1) on day 1 (D1) or after the session 1 on day 2 (D2) (Figure 5A). Figure 5B shows exemplar Arc/H1a catFISH images of CA3 regions of WT (upper row) and *Kcna2*+/− (lower row) mice for D1 (Ba and Bb), D2 (Bc and Bd). For comparison, catFISH data on retrieval of contexts on day 4 (D4) were adopted from Figure 4.

**Figure 5.**
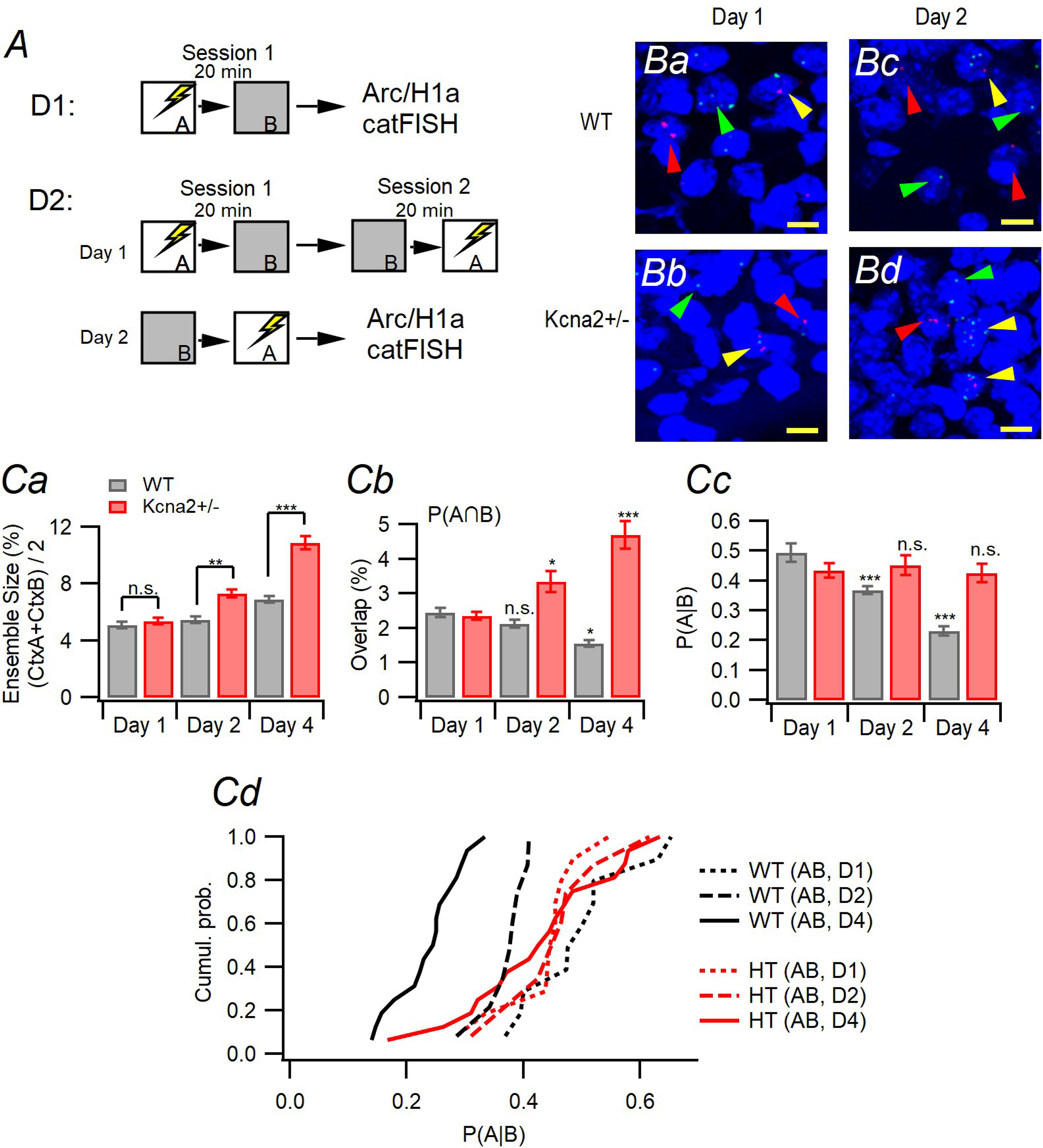
Changes in CA3 ensembles over training days of CFD task in WT and *Kcna2*+/− mice. **A**. Experimental procedures for catFISH of CA3 ensembles during training phase (Day 1 and 2), when mice do not discriminate the two contexts A and B. **B.** Representative confocal images of *H1a* and *Arc* transcripts in nuclei (*blue,* counterstained with DAPI) of the CA3 pyramidal layer from WT and Kcna2+/− mice in day 1 (*Ba*, WT; *Bb*, *Kcna2*+/−:), day 2 (*Bc*, WT; *Bd*, *Kcna2*+/−). Nuclei expressing H1a alone, Arc alone, and both are indicated by green, red, and yellow arrowheads, respectively. **Ca**. Neuronal ensemble size activated by exposure of mice to context A and B on each day. Because there was no statistical difference in the fraction of H1a(+) and Arc(+) cells in the same slice, the ensemble size was quantified as the averaged percentage of H1a(+) or Arc(+) cells among total cells in the CA3 pyramidal layer. Note that the CA3 ensemble size in *Kcna2*+/− mice (red) grew larger over training days compared to that in WT (gray). **Cb.** The fraction of CA3 cells activated in both contexts (denoted as A∩B) over training days. The fraction of A∩B cells in *Kcna2*+/− mice (red) increased over days, whereas that in WT (gray) decreased. Compared to day 1, for WT, day 2: p = 0.457, day 4: p = 0.017. For *Kcna2*+/−, day 2: p = 0.025, day 4: p < 0.001 (two-way ANOVA, simple effect analysis). **Cc.** The conditional probability for ensemble cells active in context A among those active in context B [denoted as P(A|B)]. The P(A|B) for WT mice decreased over days, but that for *Kcna2*+/− mice did not. Compared to day1, for WT, day 2: p = 0.005, day 4: p < 0.001. For *Kcna2*+/−, day 2: p = 0.678, day 4, p = 0.816 (two-way ANOVA, simple effect analysis). **Cd.** Cumulative curves for P(A|B) on each day. The overlap index [P(A|B)] in *Kcna2+/−* mice were not different from that in WT on D1. The overlap index curves of WT shifted to the left over training days, but those of *Kcna2*+/− mice did not.

Because the ensemble sizes measured from H1a and Arc in the same slice were not statistically different (probe, F_(1,136)_ = 0.893, p = 0.436; probe × genotype, F_(1, 136)_ = 0.201, p = 0.655, probe × day, F_(2,136)_ = 0.105, p = 0.900, probe × genotype × day, F_(2,136)_ = 0.380, p = 0.685; three-way ANOVA, Table 4), the ensemble size on each day was measured as the averaged number of H1a(+) plus Arc(+) cells on each slice. The CA3 ensemble size increased over training days in both genotypes, but the expansion of CA3 ensemble size in *Kcna2*+/− (red) was more pronounced compared to WT (gray) (Fig. 5Ca; genotype, F_(1,68)_ = 36.528, p < 0.001; day, F_(2,68)_ = 51.758, p < 0.001; genotype × day, F_(2, 68)_ = 12.303, p < 0.001; WT vs *Kcna2*+/− for day 1: p = 0.635, day 2: p = 0.008, day 4: p < 0.001; two-way ANOVA and simple effect analysis). Whereas in WT mice the fraction of CA3 cells active in both of contexts A and B [denoted as P(A∩B)] decreased over training days, it rather increased in *Kcna2*+/− mice (genotype, F_(1,68)_ = 38.687, p < 0.001; day, F_(2, 68)_ = 3.159; p = 0.024, genotype × day, F_(2,68)_ = 20.287, p < 0.001; two-way ANOVA; Figure 5Cb). The increase of P(A∩B) in *Kcna2*+/− mice can be attributed to the increase in the ensemble size, because the overlap index [P(A|B)] remained static over days (Figure 5Cc). In contrast, P(A|B) in WT mice decreased over days (gene, F_(1,68)_ = 10.40, p = 0.002; day, F_(2, 68)_ = 14.719; p < 0.001, genotype × day, F_(2,68)_ = 12.383, p < 0.001; two-way ANOVA). Figure 5Cd shows the changes in the cumulative distribution of the overlap index over training days for two genotypes. On day 1, the distribution in *Kcna*2+/− mice was not different from that in WT (Z = 1.118, p = 0.164; Kolmogorov–Smirnov test). The distributions in WT mice shifted to the left over training days (χ^2^ = 25.983, p < 0.001; Kruskal-Wallis test) (D1 vs. D2: Z = −2.756, p = 0.006; D2 vs. D4: Z = −3.708, p < 0.001; Kolmogorov–Smirnov test), whereas the distribution of *Kcna2*+/− mice remained unaltered (χ^2^ = 0.223, p = 0.895; Kruskal-Wallis test). In summary, the size and overlap of A and B ensembles activated by the first visit were not different between WT and *Kcna*2+/− mice. In WT mice, the A and B ensembles evolved into more distinct ones over training days. In contrast, the same training induced little pattern separation in CA3 ensembles of *Kcna2*+/− mice with the ensemble size more enlarged compared to WT.

**Table 4.** Fractions of H1a(+) and Arc(+) cells in CA3 (mean ± SEM)

| Day | Genotype | H1a (%) | Arc (%) | P(A∩B) (%) | P(A\|B) (%) |
| --- | --- | --- | --- | --- | --- |
| Day1<br>(n=10 from 3 mice) | Wildtype | 5.11 ± 0.28 | 5.07 ± 0.34 | 2.45 ± 0.14 | 49.3 ± 2.98 |
| | *Kcna2*<sup>+/-</sup> | 5.20 ± 0.31 | 5.55 ± 0.31 | 2.37 ± 0.12 | 43.5 ± 2.33 |
| Day 2<br>(n=7 from 3 mice) | Wildtype | 5.13 ± 0.35 | 5.80 ± 0.26 | 2.13 ± 0.11 | 36.9 ± 1.42 |
| | *Kcna2*<sup>+/-</sup> | 7.17 ± 0.34 | 7.43 ± 0.44 | 3.35 ± 0.30 | 45.2 ± 3.27 |
| Day 4<br>(n=16 from 4 mice) | Wildtype | 7.04 ± 0.30 | 6.57 ± 0.32 | 1.56 ± 0.10 | 23.2 ± 1.53 |
| | *Kcna2*<sup>+/-</sup> | 10.61 ± 0.66 | 11.19 ± 0.64 | 4.69 ± 0.41 | 42.6 ± 3.15 |

### Generation of CA3 region-specific hetero-knockout of Kcna2

Above results indicate that haploinsufficiency of *Kcna2* lowers the threshold for PP-LTP and impairs behavioral pattern separation (Figure 1 and 3). However, Kv1.2 subunits are expressed not only in CA3-PCs but also in other brain regions (Sheng et al., 1994; Grosse et al., 2000). To confirm that insufficiency of Kv1.2 expressed in CA3-PCs is responsible for the impairment in behavioral pattern separation, we created CA3 region-specific *Kcna2+/−* (CA3-*Kcna2+/−*) mice using Cre-lox recombination techniques (see Materials and Methods). We labelled *Kcna2* transcripts in the hippocampus of each genotype mice (postnatal week 16) using an RNAscope probe (Wang et al., 2012), and counted the number of mRNA dots in the layer 2/3 of medial entorhinal cortex (MEC) and three hippocampal regions [indicated as a (MEC), b (DG), c (CA3) and d (CA1) in Figure 6A-B]. The *Kcna2* mRNA expression level was quantified as the ratio of the number of *Kcna2* mRNA particles to the number of nuclei in the region of interest (ROI) drawn on each region. The *Kcna2* mRNA in the CA3 of *CA3-Kcna2+/−* mice (n = 5, Figure 6Bc) was lowered to c.a. 60% of that in the CA3 of *floxed-Kcna2* mice (*f-Kcna2*) (n = 5, Figure 6Ac), whereas the *Kcna2* mRNA in the MEC, DG and CA1 were not different between genotypes (Figure 6A-C; MEC, U = 7.00, p = 0.251; DG, U = 12.00, p = 0.917; CA3, U = 0.00, p = 0.009; CA1, U = 8.00, p = 0.347; Mann-Whitney U test). As a parameter for the expression level of Kv1.2, the peak amplitudes of D-type K^+^ current (I_K(D)_) were measured from CA3-PCs in *f*-*Kcna2* (Figure 6Da, left) and *CA3-Kcna2*+/− (right) mice. We regarded the slowly inactivating K^+^ outward current sensitive to 30 μM 4-aminopyridine (4-AP) as I_K(D)_ (Storm, 1988; Hyun et al. 2013, 2015). The peak amplitudes of I_K(D)_ elicited by depolarizing steps from -70 mV were significantly lower in CA3-PCs of CA3-*Kcna*2+/− mice than those of *f*-*Kcna2* mice (Figure 6D; -40 mV, black, t = 2.282, p = 0.046; -30 mV, blue, t = 2.698, p = 0.022; -20 mV, red, t = 3.885, p = 0.003; independent t-test).

**Figure 6.**
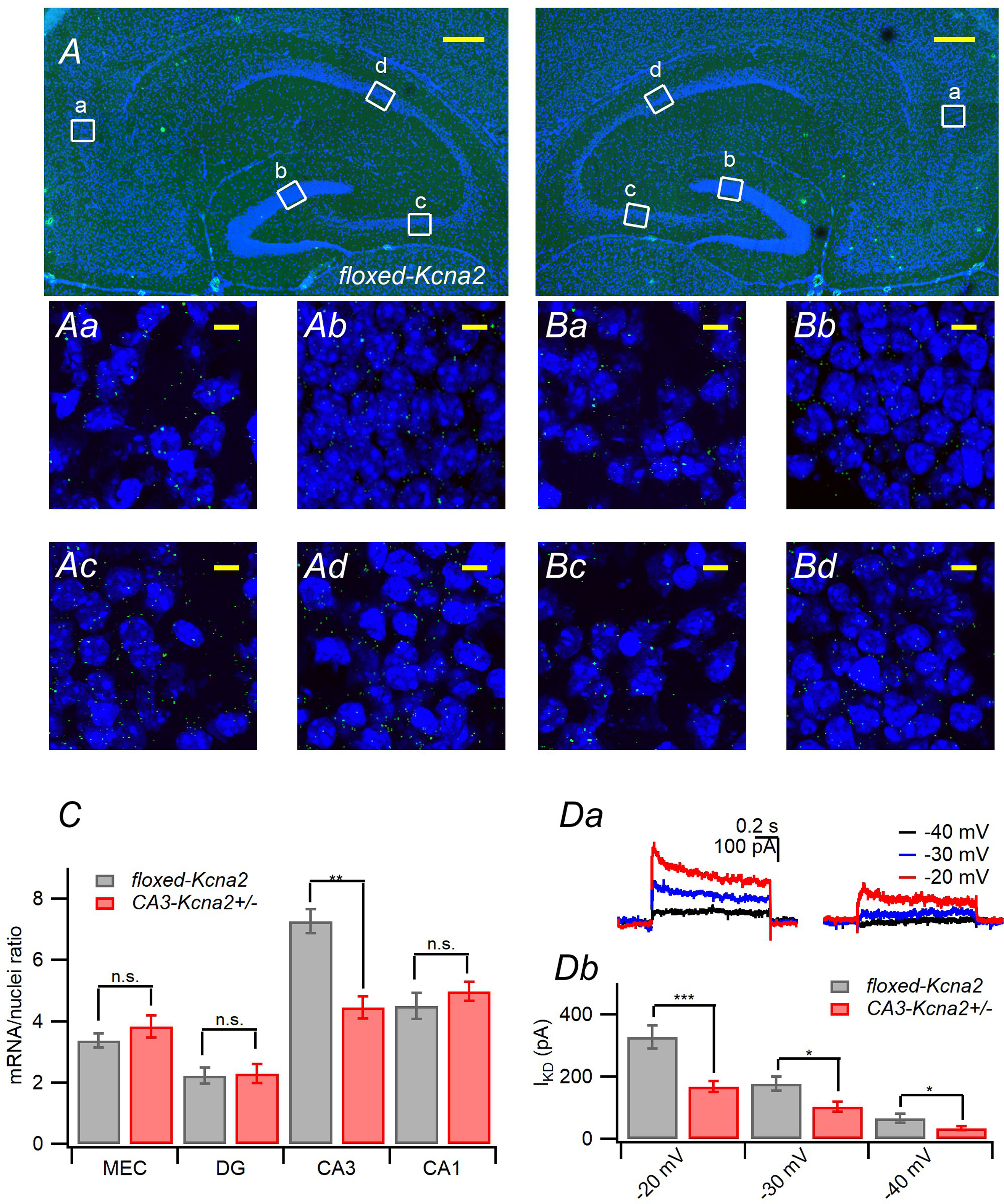
Generation of CA3 region-specific Kcna2 hetero-knockout mice. **A-B.** Confocal *Kcna2* FISH images of dorsal hippocampi from *CA3-Kcna2+/−* (A) and *floxed-Kcna2* mice (*f-Kcna2*, B). *Kcna2* transcripts were hybridized using the RNAscope probe, and visualized as green fluorescent dots. Nuclei were counterstained with DAPI (*blue*). Scale bars, 200 μm. The boxed regions [*a*, medial entorhinal cortex (MEC); *b*, DG; *c*, CA3; *d*, CA1] in each of hippocampal figures were imaged at high magnification (60 x), in which the number of red dots were counted. A part of each magnified image is shown on the below (scale bars, 10 μm). **C.** In MEC, DG and CA1, ratio of the number of *Kcna2* mRNA particles to the number of nuclei were not significantly different between *f-Kcna2* mice and *CA3-Kcna2+/−* mice (MEC, Mann-Whittney U = 7.00, p =0.251; DG, U = 12.00, p = 0.917; CA1, U = 8.00, p = 0.347, n = 5). In CA3, however, the mRNA ratio was decreased by c.a. 60% in *CA3-Kcna2 +/−* mice compared to the *f-Kcna2* mice (n = 4, U = 0.00, p = 0.009). **Da.** Representative traces for D-type K^+^ currents (*I*_K(D)_) was obtained by the arithmetical subtraction of outward potassium currents (*I*_K_) under the bath application of 30 μM 4-AP from the total *I*_K_ elicited by a depolarizing step to -20 (red), -30 (blue), and -40 (black) mV from -70 mV in the *f-Kcna2* (left) and *CA3-Kcna2*+/− (right) CA3-PCs. **Db.** Mean values for peak amplitudes of *I*_K(D)_ induced by a step depolarization to −20, −30 and −40 mV for *f-Kcna2* (gray) and *CA3-Kcna2*+/− (red) CA3-PCs.

To test if there is any alteration in the afferent inputs to CA3-PCs of *CA3-Kcna2*+/− mice, we assessed short-term synaptic plasticity of MF and PP-EPSCs, because reduction of axonal Kv1 channels is expected to broaden the axonal APs and increase in synaptic release probability to lower paired pulse ratio of EPSCs (Kole et al., 2007). Previously, it was noted that PP expresses both of Kv1.1 and Kv1.2, while MFs express primarily Kv1.1 (Monaghan et al. 2001). Consistently, short-term facilitation (STF) of PP-EPSCs evoked by 5 pulses at 20 or 50 Hz was reduced in *Kcna1*+/− and *Kcna2*+/− mice, but not *CA3-Kcna2*+/− mice. For MF-EPSCs evoked by 3 pulses at 50 Hz, STF was reduced only in *Kcna1*+/ mice (Table 5, Figure 6-figure supplement 1).

**Table 5.**
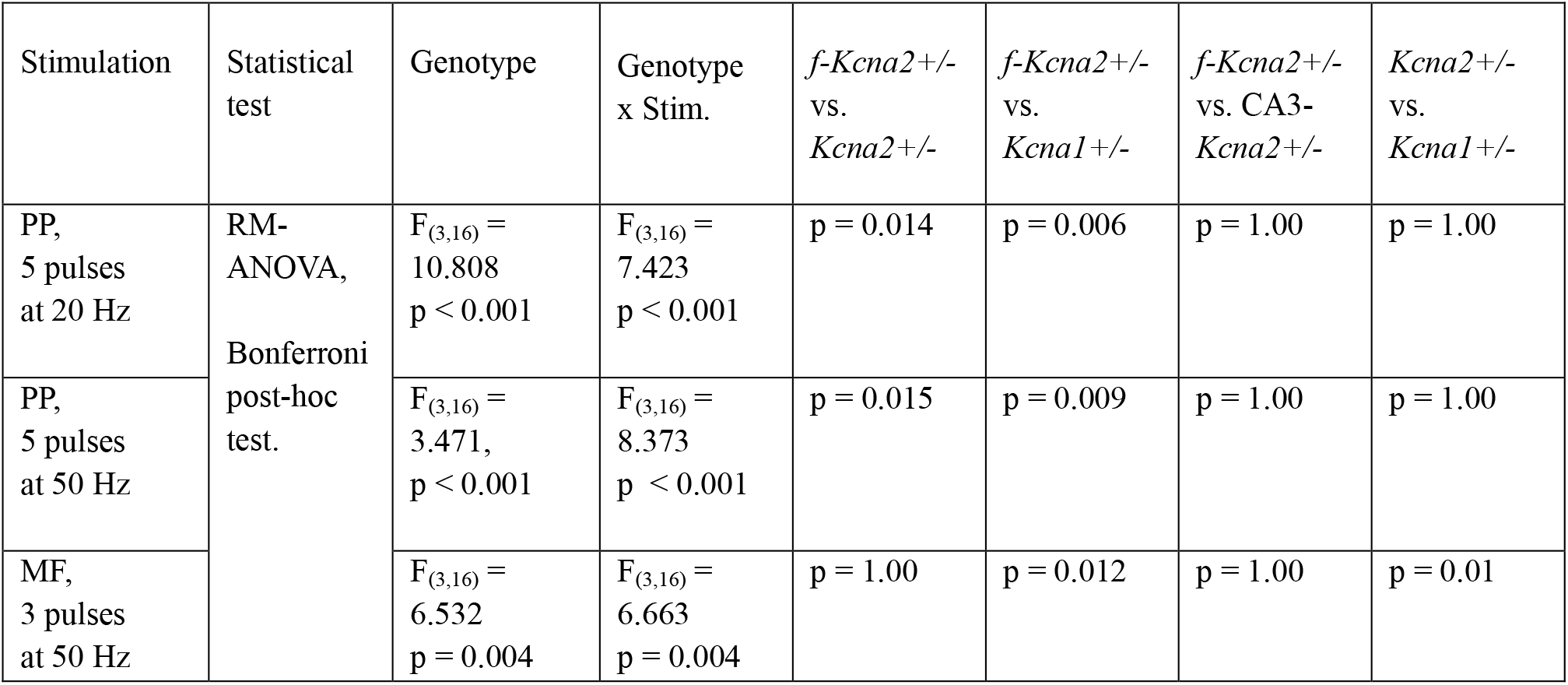
Short term plasticity at PP and MF synapses of CA3-PCs

| Stimulation | Statistical test | Genotype | Genotype x Stim. | *f-Kcna2*<sup>+/-</sup> vs. *Kcna2*<sup>+/-</sup> | *f-Kcna2*<sup>+/-</sup> vs. *Kcna1*<sup>+/-</sup> | *f-Kcna2*<sup>+/-</sup> vs. CA3-*Kcna2*<sup>+/-</sup> | *Kcna2*<sup>+/-</sup> vs. *Kcna1*<sup>+/-</sup> |
| --- | --- | --- | --- | --- | --- | --- | --- |
| PP, 5 pulses at 20 Hz | RM-ANOVA, Bonferroni post-hoc test. | F<sub>(3,16)</sub> = 10.808<br>p < 0.001 | F<sub>(3,16)</sub> = 7.423<br>p < 0.001 | p = 0.014 | p = 0.006 | p = 1.00 | p = 1.00 |
| PP, 5 pulses at 50 Hz |  | F<sub>(3,16)</sub> = 3.471,<br>p < 0.001 | F<sub>(3,16)</sub> = 8.373<br>p < 0.001 | p = 0.015 | p = 0.009 | p = 1.00 | p = 1.00 |
| MF, 3 pulses at 50 Hz |  | F<sub>(3,16)</sub> = 6.532<br>p = 0.004 | F<sub>(3,16)</sub> = 6.663<br>p = 0.004 | p = 1.00 | p = 0.012 | p = 1.00 | p = 0.01 |

Next, we examined MF-induced LTP-IE of CA3-PCs in CA3-*Kcna2+/−* mice and their WT littermates (*f-Kcna2*) in the third postnatal week because Cre/loxP recombination is detectable on postnatal day 14 in the CA3 region (Nakazawa et al., 2002). In addition, we previously reported that the degree of LTP-IE is not different between the third and 12th postnatal week mice (Eom et al., 2019). LTP-IE was assessed by input conductance (G_in_) and first spike latency, which were measured as in Figure 1B. In *f-Kcna2* CA3-PCs, 20 Hz MF stimulation reduced G_in_ (75.3 ± 2.7%, n=6, t = 7.714, p = 0.001, paired t-test, Figure 7Aa-Ab) and first spike latency (t = 3.855, p = 0.012, n = 6, paired t-test, Figure 7Ac). In contrast, MF-induced LTP-IE was completely abolished at CA3-PCs of CA3-*Kcna2+/−* (Figure 7A; G_in_, 99.07 ± 2.8%, t = 0.541, p = 0.612; AP latency, n = 6, t = −0.947, p = 0.397, paired t-test). The baseline G_in_ and AP latency in CA3-*Kcna2+/−* CA3-PCs were lower than those of *f-Kcna2*, and was not significantly different from those of *Kcna2+/−* mice (Figure 7Ab-Ac). Next, we examined MF-induced heterosynaptic potentiation of PP-EPSPs for the CA3-*Kcna2+/−* and *f-Kcna2* mice. LTP of PP-EPSP was readily induced in the *f-Kcna2* CA3-PCs (185.46 ± 2.59%, n = 5, t = −5.874, p = 0.004; paired t-test), but it was abolished in the *CA3-Kcna2+/−* (91.30 ± 11.91%, n = 6, t = 0.348, p = 0.742; paired t-test; Figure 7Ba and 7Bb). Supporting the notion that LTP-IE underlies the heterosynaptic LTP of PP-EPSPs (Hyun et al., 2015), 20 Hz MF stimulation did not alter PP-EPSCs (Figure 7Bc).

**Figure 7.**
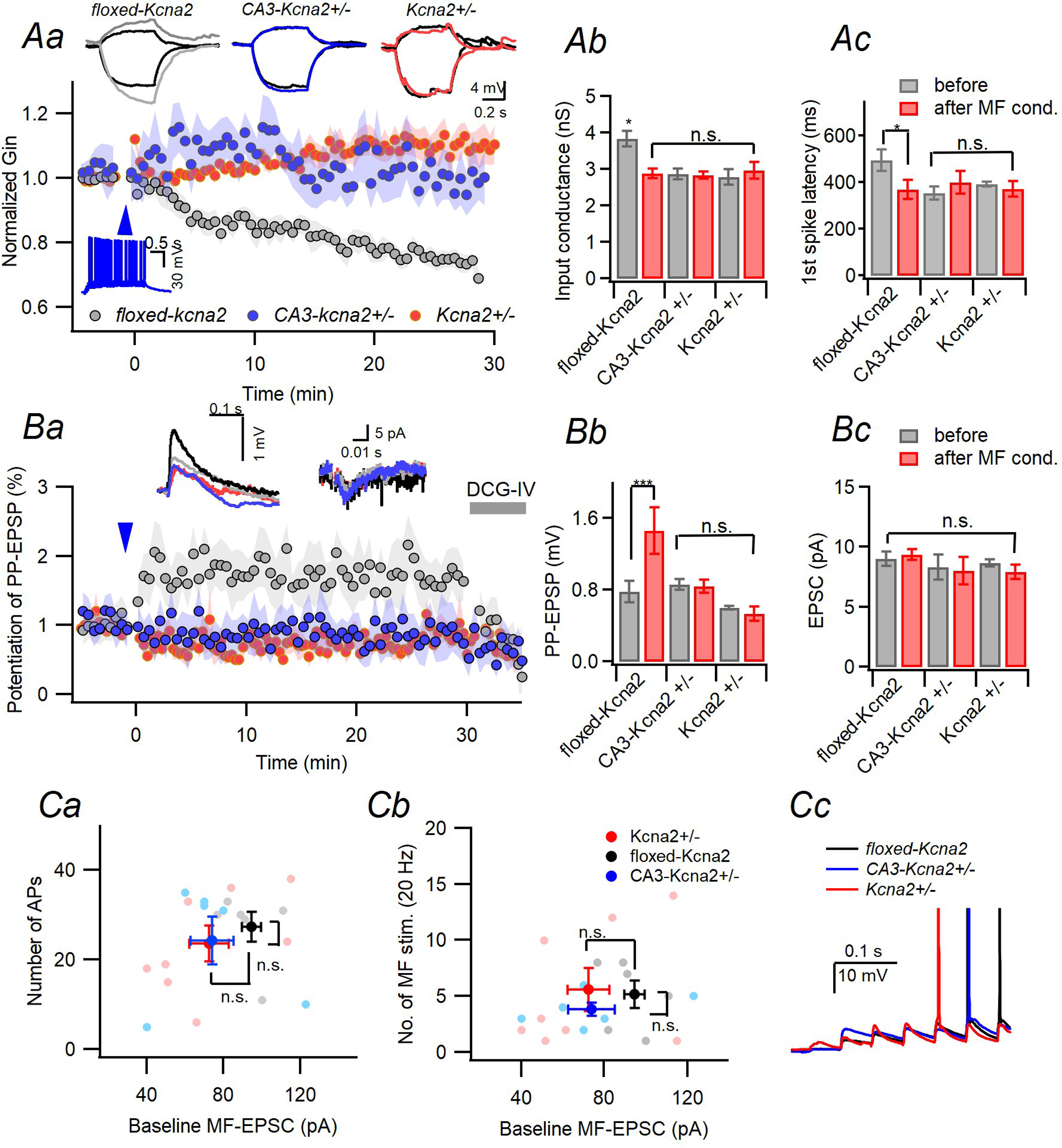
CA3 region-specific *Kcna2* +/− mice lacks MF-induced LTP-IE. **Aa.** MF conditioning (20 Hz MF stimulation, *blue arrowhead*) induced a decrease in input conductance (G_in_) in *f-Kcna2* (*gray*), but not in *CA3-Kcna2+/−* (*blue*) and *Kcna2+/−* (*red*) CA3-PCs. *Insets*: Representative traces for postsynaptic response to 20 Hz MF stimulation (*blue trace in the main figure*) and voltage responses to subthreshold current injections (+10/-30 pA, *upper*). **Ab-Ac.** Mean values for baseline (*gray*) and post-conditioning (*red*) input conductance (*Ab*) and first AP latency (*Ac*). G_in_ and first AP latency were measured in the same way as in Figure 1B. The baseline G_in_ and first AP latency of *CA3-Kcna2+/−* CA3-PCs (n=6) were lower than that of *f-Kcna2* (n=6), and the same as that of *Kcna2+/−* (n=14). (For G_in_, F_(2,24)_ = 6.036, p = 0.008; f-*Kcna2 vs.* CA3-*Kcna2+/−*, p = 0.045; *f-Kcna2* vs. *Kcna2+/−*, p = 0.008; CA3-*Kcna2+/−* vs. *Kcna2+/−*, p = 1.00; For AP latency, F_(2,28)_ =5.714, p = 0.017; *f-Kcna2* vs*. Kcna2+/−*, p = 0.112; CA3-*Kcna2+/−* vs. *f-Kcna2*, p = 0.019; CA3-*kcna2+/−* vs. *Kcna2+/−*, p = 1.00; one-way ANOVA and Bonferroni post-hoc test). **Ba.** MF conditioning-induced heterosynaptic potentiation of PP-EPSPs in *f-Kcna2* (*gray*), *CA3-Kcna2+/−* (*blue*), and *Kcna2+/−* (*red*) CA3-PCs. *Insets*: Representative traces for PP-EPSPs (*left*) and PP-EPSCs (*right*) before (*black*) and after MF conditioning. The color codes for post-conditioning traces are same as the main figure. **Bb.** Amplitude of PP-EPSPs before (*gray*) and after (*red*) MF conditioning. MF conditioning induced heterosynaptic potentiation of PP-EPSP in *f-Kcna2* (n=5), but not in *CA3-Kcna2+/−* (n=6) and *Kcna2*+/− (n=8) CA3-PCs. **Bc.** Mean amplitude of PP-EPSCs was not altered before and after MF conditioning (*f-Kcna2*, t = −0.451, p = 0.675; CA3-*Kcna2*+/−, t = 0.500, p = 0.638; *Kcna2+/−*, t = 0.518, p = 0.632, paired t-test). **Ca** and **Cb.** The number of APs elicited by 20 Hz / 2 s MF stimulation (*Ca*) and MF stimulation required to elicit the 1st AP (*Cb*) as a function of baseline MF-EPSC amplitude for *f-Kcna2* (black), *CA3-Kcna2*+/− (blue) and *Kcna2*+/− (red). No statistical difference was found between genotypes in AP numbers (*Ca*, F_(2,19)_ = 0.294, p = 0.749, one-way ANOVA) and MF stimulation numbers (*Cb*, F_(2,19)_ = 0.864, p = 0.439, one-way ANOVA). **Cc.** Representative EPSP traces evoked by 20 Hz MF stimulation during which the first AP was evoked in the three genotype CA3-PCs.

Previously we demonstrated that downregulation of Kv1.2 after induction of LTP-IE enhances specifically PP-EPSPs with little effect on MF-EPSPs (Hyun et al., 2015). To test if a reduction of Kv1.2 channels has any effect on E-S coupling of synaptic inputs to proximal dendrites, the number of APs elicited during 20 Hz MF stimulation for 2 s (Figure 7Ca) and the number of MF stimulation required to elicit the 1^st^ AP in the postsynaptic CA3-PC (Figure 7Cb) was plotted as a function of baseline MF-EPSC in the three genotype mice (*f-Kcna2*, *Kcna2*+/−, *CA3-Kcna2+/−*). Both parameters were not different between the three genotypes, confirming that reduction of Kv1.2 has little effect on the proximal dendritic excitability.

### Insufficiency of Kv1.2 in CA3-PCs is responsible for the impaired pattern separation

To test whether haploinsufficiency of *Kcna2* confined to CA3-PCs is sufficient to impair the CFD task, we repeated the behavioural test of Figure 3A for *f-Kcna2* and *CA3-Kcna2+/−* mice. In *CA3-Kcna2*+/− mice the expression of Cre recombinase is driven by the promoter of a kainate receptor subunit, KA-1, which is strongly expressed in CA3. KA1, however, is expressed in 10% of dentate and cerebellar granule cells too (Nakazawa et al., 2002). To rule out the possibility that reduction of Kv1.2 in brain regions other than CA3 contributes to behavioural phenotype, we delivered AAV encoding Cre-mCherry or GFP under the CaMKIIα promoter (AAVcre and AAVgfp, respectively) to dorsal and ventral CA3 regions of *f-Kcna2* mice through stereotaxic techniques (see materials and methods). The AAV-injected mice were subject to behavioral test and post-hoc *ex vivo* experiments 4 weeks after the viral injection.

On day 4-5, all genotype and AAV-injected mice showed similar kinetics of freezing in context A (Figure 8Aa, Table 6), and no difference was observed in freezing behavior among genotypes and AAV-injected mice [WT (n=11), *f-Kcna2* (n=4), *CA3-Kcna2*+/− (n=4), *Kcna2*+/− (n=9) mice, and AAVcre (n=4) or AAVgfp (n=4)-injected mice] and between contexts A and B (Figure 8Ab, Table 6), indicating that generalization between the two contexts is similar in the three genotype mice. On the subsequent 9 days (day 6 -14, phase 3), mice daily visited both contexts with receiving a foot shock only in context A. The discrimination ratio of *f-Kcna2* and AAVgfp-injected mice, but not *CA3*-*Kcna2+/−* and AAVcre-injected mice, began to rise already on day 7 (Figure 8Ba). During acquisition of the discrimination over the 9 training days, *f-Kcna2* and AAVgfp-injected mice quickly discriminated context B from context A similar to WT mice, whereas CA3*-Kcna2+/−*, *Kcna2*+/− and AAVcre-injected mice showed significant delay for discrimination (Figure 8Ba, Table 6). Notably, the learning curves of *CA3-Kcna2+/−* were not different from that of *Kcna2*+/− mice (p = 1.00) and AAVcre-injected mice (p = 1.00), whereas it was different from those of WT (p < 0.001), *f-Kcna2* (p = 0.02), and AAVgfp-injected mice (p = 0.005; RM-ANOVA Bonferroni post-hoc test, Figure 8Ba). The discrimination was impaired in the *CA3-Kcna2+/−* and AAVcre-injected mice on day 9 (Figure 8Bb; comparing with WT, *CA3-Kcna2+/−*: p < 0.001, *Kcna2*+/−: p < 0.001, *f-Kcna2*: p = 0.680, AAVgfp: p = 0.466, AAVcre: p < 0.001, two-way ANOVA), but it was eventually acquired on day 14, similar to WT and *f-Kcna2* (Figure 6Bc, Table 6). After the behavioral test, AAV-injected mice were sacrificed and subject to *ex vivo* experiments to examine the AAV infection area and reduction of *I*_K(D)_ in CA3-PCs (Figure 8C-E). Confirming that mCherry or GFP signal is confined to the CA3 region (Figure 8C-D), *I*_K(D)_ was measured from GFP(+) and mCherry(+) CA3-PCs using the same methods as in Figure 6D (Figure 8E). The peak amplitude of *I*_K(D)_ activated by a depolarizing step to -20 mV from -70 mV in CA3-PCs of AAVcre-injected mice (Figure 8Eb) was significantly lower than that of AAVgfp-injected mice (Figure 8Ea; AAVgfp, 325.7 ± 30.65 pA; AAVcre, 168.74 ± 25.73 pA, t = 3.921, p = 0.004, independent t-test, Figure 8Ec).

**Table 6.** Statistical test for experiments shown in Figure 8

| Experiments | Statistical test | 1<sup>st</sup> variable<br>(Genotype) | 2<sup>nd</sup> variable<br>(Context / Time /<br>Day) | Interaction |
| --- | --- | --- | --- | --- |
| Figure 8Aa | RM- ANOVA | $F_{(5,30)} = 0.616$<br>$p = 0.689$ | Time,<br>$F_{(9,270)} = 9.799$<br>$p < 0.001$ | Geno x Time,<br>$F_{(9,270)} = 0.523$<br>$p = 0.996$ |
| Figure 8Ab<br>(Day 4-5) | two-way ANOVA | $F_{(5,67)} = 2.336$<br>$p = 0.054$ | Context<br>$F_{(1,67)} = 0.334$<br>$p = 0.566$ | Geno x Context.<br>$F_{(5,67)} = 0.226$<br>$p = 0.950$ |
| Figure 8Ba<br>(Discrimination<br>ratio) | RM- ANOVA<br>Bonferroni post-hoc test | $F_{(6,39)} =$<br>11.614<br>$p < 0.001$ | Day<br>$F_{(8,312)} = 27.433$<br>$p < 0.001$ | Geno x Day,<br>$F_{(48,312)} = 2.930$<br>$p < 0.001$ |
| Figure 8Bb<br>(Day 9) | two-way<br>ANOVA,<br>simple effect<br>analysis | $F_{(5,72)} = 13.408$<br>$p < 0.001$ | Context<br>$F_{(1,72)} = 47.450$<br>$p < 0.001$ | Geno x Context,<br>$F_{(5,72)} = 6.269$<br>$p < 0.001$ |
| Figure 8Bc<br>(Day 14) | two-way<br>ANOVA, | $F_{(5,72)} = 3.529$<br>$p = 0.007$ | Context<br>$F_{(5,72)} = 438.86$<br>$p < 0.001$ | Geno x Context.<br>$F_{(5,72)} = 1.520$<br>$p = 0.197$ |

Altogether, despite that release probability at PP-CA3 synapses is enhanced both in *Kcna1*+/− and *Kcna2*+/− mice, *Kcna1*+/− mice was normal in CFD (Figure 3) and in pattern separation of CA3 ensembles (Figure 4), suggesting that impaired pattern separation in *Kcna2*+/− mice cannot be attributed to presynaptic Kv1.2 haploinsufficiency. Because both *CA3-Kcna2*+/− mice and AAVcre-injected mice displayed impairments in the CFD task, it is unlikely that insufficiency of Kv1.2 in brain regions other than hippocampal CA3 is responsible for the impairment in CFD. Given that performance in CFD task is closely correlated with pattern separation in CA3 ensembles, insufficiency of Kv1.2 in CA3-PCs rather than other brain regions might be responsible for the impaired pattern separation of CA3 ensembles in the *Kcna2*+/− mice.

**Figure 8.**
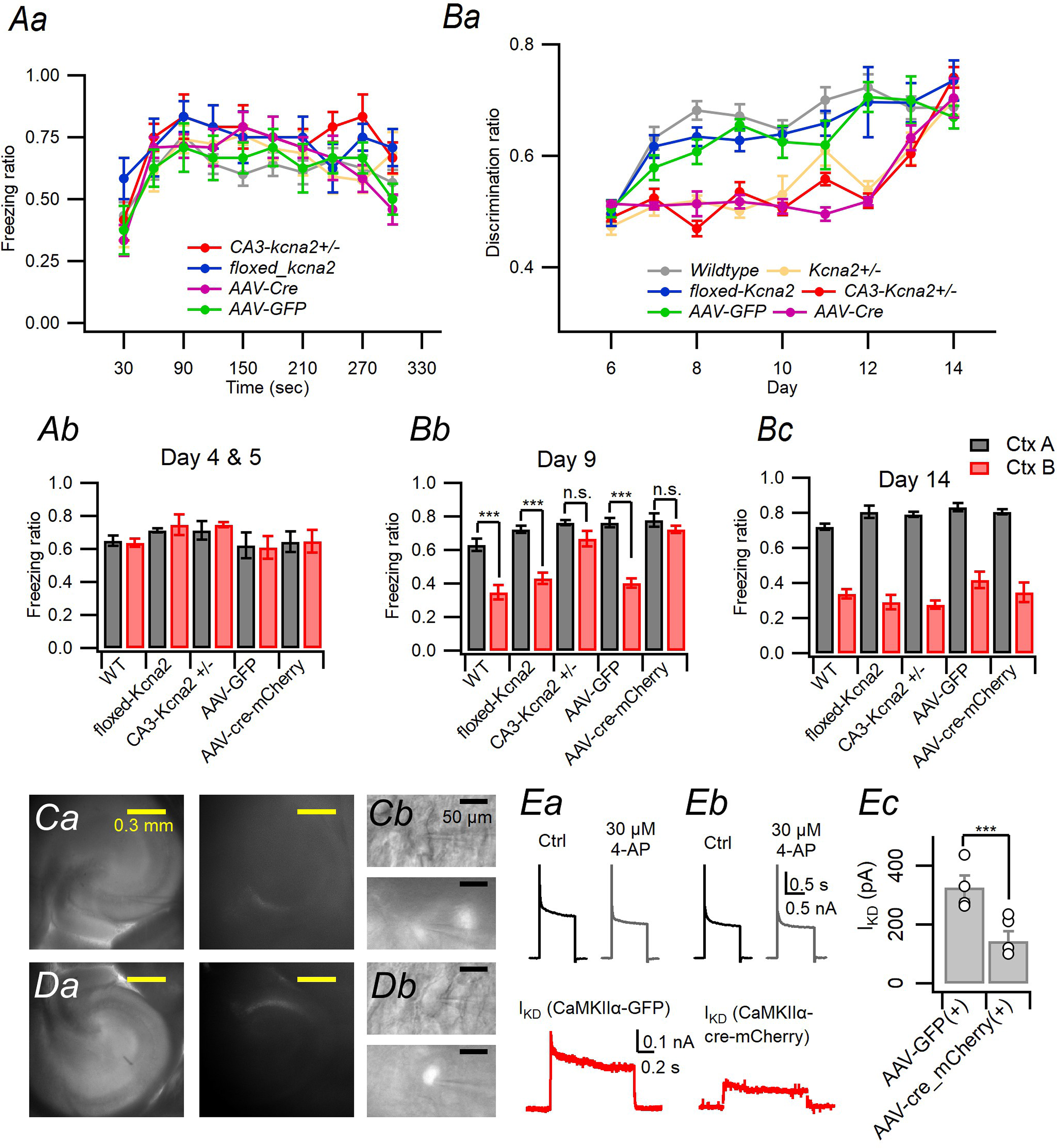
Kv1.2 expressed in CA3-PCs is essential for rapid contextual discrimination. **Aa.** Kinetics of freezing behaviour over 5 min in context A on day 4-5. **Ab.** On day 4 and 5, all genotype mice could not discriminate the two contexts A and B. **Ba.** WT, *f-Kcna2* and AAVgfp-injected mice showed significantly earlier rise in the discrimination ratio than *CA3-Kcna2+/−*, *Kcna2+/−* and, AAVcre-injected mice over the day 6 to 14 (WT vs. *f*-*Kcna2*, p = 1.00; WT vs. AAV-GFP injected, p = 1.00; WT vs. *CA3-Kcna2+/−*, p < 0.001; WT vs. *Kcna2*+/−, p < 0.001; WT vs. AAVcre-injected: p < 0.001; *f-Kcna2* vs. CA3-*Kcna2+/−* p < 0.012; CA3-*Kcna2+/−* vs *Kcna2*+/−, p = 1.00; AAVgfp vs. AAVcre: p = 0.036; RM-ANOVA and Bonferroni post hoc test). **Bb.** On day 9, WT, *f-Kcna2* and AAVgfp-injected mice discriminated the two similar contexts, whereas CA3-*Kcna2+/−* and AAVcre-injected mice still could not. Comparing the freezing ratio for A vs B in each genotype, WT: p < 0.001, *f-Kcna2*: p < 0.001, CA3-*Kcna2*+/−: p = 0.185, *Kcna2*+/−: p = 0.899, AAVcre: p = 0.446, AAVgfp: p < 0.001 (two-way ANOVA, simple effect analysis). **Bc**. On day 14, initial discrimination deficit for *CA3-Kcna2+/−* was rescued by repetitive training. **C-D**, IR-DIC (*Ca* and *Da*, left) and epifluorescence images (right, excited by 488 or 588 nm for GFP or mCherry, respectively) of the hippocampi from the AAVcre-injected (*Ca*) or AAVgrp-injected (*Da*) mice. Corresponding magnified IR-DIC (*Cb* and *Db*, upper) and epifluorescence images (lower). **Ea-b**, *I*_K(D)_ in GFP(+) and mCherry(+) CA3-PCs from acute slices prepared from AAVgfp-injected and AAVcre-injected mice, respectively. *I*_K(D)_ (lower, red) was obtained from difference in the K^+^ outward current activated by a depolarizing step to -20 mV from -70 mV before (upper left, black) and after the bath application of 30 μM 4-AP (upper right, gray) in CA3-PCs of AAVgfp-injected (*Ea*) and AAVcre-injected (*Eb*) mice. **Ec**, Mean values for peak amplitudes of *I*_K(D)_.

## Discussion

Despite extensive studies on the synaptic plasticity in the CA3 region (reviewed in Rebola et al., 2017), the cognitive and behavioral consequences of altered synaptic plasticity in the CA3 network largely remains to be elucidated. For pattern separation, studies on its cellular mechanisms have mainly focused on DG (McHugh et al., 2007; Nakashiba et al., 2012; KimKR et al., 2020), but little understood in the CA3 region. In the present study, we found that insufficiency of Kv1.2 subunits in CA3-PCs enhances E-S coupling of PP synaptic inputs to lower the PP-LTP threshold (Figure 1 and 2), and closely correlates with deficit in rapid contextual discrimination (Figure 3 and 8) and with impaired pattern separation of CA3 ensembles activated by retrieval of two slightly different contexts (Figure 4).

The size and overlap of CA3 ensembles activated by the first visit to contexts A and B on day 1 were not different between WT and *Kcna2*+/− mice, but the two ensemble parameters diverged over training days (Figure 5). The *Kcna2*+/− mice displayed rapid enlargement of the CA3 ensemble size with the overlap index static over training days, while WT mice displayed relatively stable ensemble size with decreasing overlap index. <u>Because the ensemble properties of *Kcna2*+/− mice were initially similar to WT, the abnormal evolvement of CA3 ensembles in *Kcna2*+/− mice cannot be explained by increased dendritic excitability alone, but implies that the CA3 networks of *Kcna2*+/− mice undergo plastic changes different from that of WT mice over subsequent training days</u>. Given that the PP-LTP threshold is lowered by Kv1.2 haploinsufficiency, PP-LTP seems to be one of major players in the evolvement of CA3 ensembles during the training phase. Therefore, these results support the view that Kv1.2-dependent regulation of PP-LTP threshold may be crucial for stabilizing the ensemble size and pattern separation. It was previously shown that hippocampal learning induces growth of filopodia from MF terminals, which increases feedforward inhibition (FFI) triggered by MF inputs, and that the CA3 ensemble size is abnormally enlarged by contextual fear conditioning in the mutant mice in which the filopodial growth is abolished (Ruediger et al., 2011). This study together with our results suggest that balanced plastic changes at excitatory and inhibitory synapses are essential for the CA3 ensemble size to be maintained stable during hippocampal learning, and this may be the case for the wildtype mice. On the other hand, as *Kcna2*+/− mice are repeatedly exposed to the dangerous context, CA3 network may undergo too promiscuous plastic changes at PP synapses to be counterbalanced by plastic changes at inhibitory synapses such as new filopodial growth from MF terminals.

Previous studies indicate that Kv1.1 subunits are preferentially expressed in the soma and axonal compartments of CA3-PCs (Monaghan et al., 2001; Rama et al., 2017), whereas Kv1.2 subunits are in both of somatodendritic and axonal compartments (Sheng et al., 1994; Wang et al., 1994; Hyun et al., 2015). Moreover, PP expresses both Kv1.1 and Kv1.2, while MFs express Kv1.1 but not Kv1.2 (Monaghan et al., 2001). Consistently, both of *Kcna1*+/− and *Kcna2*+/− CA3-PCs displayed similar increases in the intrinsic excitability of CA3-PCs (Figure 1B) and similar changes in short-term plasticity at PP synapses (Figure 6-figure supplement 1). Nevertheless, the PP-LTP threshold (Figure 1D), behavioral phenotype (Figure 3), and the CA3 ensemble overlap (Figure 4D) were not altered in *Kcna1*+/− mice (Figure 1D, 3 and 4D). These results suggest that larger ensemble size in *Kcna2+/−* cannot be attributed to insufficiency of *Kcna2* in the axonal compartment of afferent fibers but caused by subunits located in somatodendritic compartment. Supporting this view, performance in CFD task of *CA3-Kcna2+/−* or AAVcre-injected mice were impaired similar to *Kcna2+/−* mice (Figure 8B).

### PP-LTP threshold and CA3 ensemble size

Our results imply that lowered PP-LTP threshold correlates with abnormal enlargement of the ensemble size during repeated exposure of the contexts. Given that PP synaptic inputs that were already strengthened during a previous episode contribute to the ensemble size on retrieval, sparse encoding at PP synapses would be essential for keeping the ensemble size stable. <u>MF inputs are expected to activate a small number of CA3-PCs because of low convergence of MFs to CA3-PC together with sparse firing of dentate GCs. When PP-LTP depends on concurrent MF inputs or MF-dependent priming, the sparse MF inputs to CA3 may ensure sparse encoding of memories at PP-CA3 synapses.</u> In contrast to MFs, PP makes densely distributed synapses on CA3-PCs (Amaral et al, 1990), and thus nonspecific lowering of PP-LTP threshold caused by *Kcna2* haploinsufficiency is expected to have more profound effects on the CA3 ensemble size than lowering the threshold by MF-dependent priming. Therefore, Kv1.2 in CA3-PCs seems to <u>reduce the probability for PP inputs alone to induce LTP without a help of MF inputs</u> by keeping the LTP threshold high, and plays a crucial in limiting the expansion of CA3 neuronal ensemble size during hippocampal learning.

### MF-dependent encoding at PP synapses and pattern separation of CA3 ensembles

Because MF inputs convey a decorrelated version of EC ensemble pattern to CA3 and dominate firing of CA3-PCs during a learning phase, limiting the LTP induction at PP synapses in CA3-PCs receiving strong MF inputs is thought to be essential for formation of sparse and discrete ensembles in CA3, and thus pattern separation. In *Kcna2*+/− mice, however, non-specific lowering of PP-LTP threshold renders the PP-LTP induction less dependent of MF inputs, and thus the overlap between ensembles in the input layer (EC) may be transferred to the CA3 network without decorrelation. This view may explain why the overlap of CA3 ensembles in *Kcna2+/−* mice is larger than that expected from their large ensemble size (Figure 4Da-Db). On the other hand, the pattern completion behavior probed by PECFC was not altered in *Kcna2+/−* mice (Figure 3D). Given that pattern completion is mediated by A/C fibers, this result is consistent with our previous report that downregulation of Kv1.2 does not alter EPSPs evoked by A/C fibers (Hyun et al., 2015). Taken together, an increased probability for MF-independent PP-LTP may render the *Kcna2*+/− mice impaired in formation of discrete representation of memories for two slightly different contexts during learning phase. Such impaired decorrelation of memories may result in generalization of the two contexts by a pattern completion process.

### Possible role of MF-induced metaplasticity at PP-CA3 synapses in the CA3 network computation

We have previously shown that repetitive MF inputs to a CA3-PC downregulate Kv1.2 subunits in Ca^2+^- and Zn^2+^-dependent manner, resulting in LTP-IE and specific potentiation of PP-EPSPs (Hyun et al, 2013, 2015; Eom et al., 2019). LTP-IE, which involves global excitability changes over the distal apical dendritic arbor, is unlikely to directly mediate memory encoding. Therefore, the role of LTP-IE in memory encoding should be studied with respect to the plastic changes of PP-EPSCs, which reflects synapse-specific events. The present study showed that LTP-IE greatly lowers the threshold of PP synaptic inputs required for subsequent plastic changes of PP-EPSCs (Figure 2), indicating that not only concurrent MF inputs but also previous history of MF inputs may facilitate encoding at PP-CA3 synapses.

Lowering PP-LTP threshold by MF-induced metaplasticity may potentially increase the size and overlap of CA3 ensembles, and thus may have negative effects on pattern separation. The CA3 ensemble size in *Kcan2*+/− mice was 1.5 times larger on day 4 (Figure 4D), even though *Kcna2+/−* CA3-PCs lack MF-induced metaplasticity, suggesting that non-specific lowering of PP-LTP threshold caused by *Kcna2* haploinsufficiency seems to exert more adverse effects on pattern separation compared to MF-induced metaplasticity. Although the CA3 neuronal ensemble size should be larger in the presence of MF-induced metaplasticity than without it, the increment may be very limited, because induction of LTP-IE requires <u>high frequency MF inputs</u> (Eom et al., 2019). To figure out the role of MF-induced metaplasticity in pattern separation and CA3 network computation, it needs to be elucidated the cellular mechanisms underlying depotentiation of LTP-IE, because the role of MF-induced metaplasticity in CA3 network computation would heavily depend on how long the primed state of CA3-PCs is maintained *in vivo*. Although MF-induced LTP-IE, which corresponds to the metaplastic or primed state, lasts longer than 30 min *in vitro* (Hyun et al., 2015; Eom, 2019; Figure 7A), it may not be the case *in vivo*. If the primed state of CA3-PCs is maintained until an animal experiences the next episode, which activates the second ensembles, such CA3-PCs would be re-activated by diffusely incoming PP inputs during the 2nd episode independent of MF inputs. This scenario raises a possibility that a subset of CA3-PCs primed during the 1st episode may play a key role in bridging the two ensembles encoding the 1^st^ and 2^nd^ episodes even if the two episodes are temporally discontiguous. In this case, MF-induced metaplasticity should have more or less adverse effects on pattern separation because the primed subset will be activated by both contexts associated with the two episodes. If metaplastic states of CA3-PCs are relatively short lasting, however, it may simply subserve MF-dependent encoding at PP synapses of a single episodic memory, and thus may have little effect on pattern separation.

## Materials and Methods

### Animals and ethical approval

All of studies and experimental protocols described in this article were approved by the Institutional Animal Care and Use Committee (IACUC) of Seoul National University. The animals were maintained in standard environmental conditions (Temperature: 25 ± 2 °C, Humidity: 60 ± 5 %, Dark/Light cycle: 12/12h) and monitored under veterinary supervision by Institute for Experimental Animals, Seoul National University College of Medicine.

### Kv1.1 or Kv1.2 mutant mice

Heterozygous *Kcna1* and *Kcna2* knock-out mice (C3HeB/FeJ-*Kcna1*^tm1tem^ and C3HeB/FeJ-*Kcna2*^tm1tem^, denoted as *Kcna1*+/− and *Kcna2*+/−, respectively) were kindly donated by Dr. Bruce Tempel (University Washington, Seattle) and purchased from The Jackson Laboratory (BarHarbor, ME, USA; Donating Investigator: Dr. Bruce Tempel, University of Washington School of Medicine), respectively. These mice were maintained under the veterinary supervision by Institute for Experimental Animals, Seoul National University College of Medicine. By inter-crossing heterozygote mice, we bred homozygous knock-out mice, heterozygote mice and wild-type littermate (WT) for two genotypes. For genotyping, DNA was isolated from the tail of each mouse in a litter aged 6–8 days as described by (Brew et al., 2007). Detailed protocols are available online (Kcna2: https://www.jax.org/Protocol?stockNumber=010744&protocolID=24908; Kcna1: https://www.jax.org/Protocol?stockNumber=003532&protocolID=27668). Although Kcna2−/− and Kcna1−/− mice had a severely reduced life span (range P18–P23; P4-6 wks, respectively) (Brew et al., 2007), they appeared normal during the first 2 weeks of their life.

### Generation of CA3 region-specific Kv1.2 mutant mice

Generation of mice bearing the ‘Knockout-first’ allele of *Kcna2* <Kcna2tm1a(EUCOMM)Wtsi> was performed by Macrogen Inc. (Seoul, Republic of Korea). JM8A3.N1 ES cells (Clone ID: EPD0544_3_G05; Allele name: Kcna2tm1a(EUCOMM)Wtsi; Coat color: agouti) from C57BL/6N mice were imported from the EUCOMM (European Conditional Mouse Mutagenesis; Helmholtz Zentrum, München, Neuherberg, Germany) Project. The JM8A3.NA ES cell was cultured to acquire enough number of ES cells to injection to C57BL/6N blastocysts to form new chimeric embryos. The embryos were transferred into the uteri of pseudopregnant recipient ICR female mice (5 blastocysts per conus uteri). Acquired male chimeric mice (F_0_) were bred with C57BL/6N wildtype females to transfer Kcna2tm1a(EUCOMM)Wtsi allele to germ cells. Germ-line transmission of the allele was analyzed by PCR in a F1 litter aged 6–8 days. After the screening of Kcna2tm1a(EUCOMM)Wtsi from F1 offspring mice (hereafter ‘tm1a’ mice), they were mated with general “FRT deleter” mice (Protamine-FLPe) to cut of both the LacZ and neomycin site, leading to the generation of ‘*floxed-Kcna2*’ mice. A CA3-specific cre-expressing mice (C57BL/6-Tg(Grik4-cre)G32-4Stl/J) was mated with tm1c mice to cut exon 3 via Cre-lox recombination, leading to deletion of *Kcna2* in CA3 subregion (hereafter called ‘CA3-*Kcna2+/−*’).

### Stereotaxic surgery and intrahippocampal injection of adeno-associated virus

A pair of adeno-associated virus (AAV) was kindly provided by Dr. Joung-Hun Kim (Dept. of Life sciences, POSTECH) (AAV-CaMKIIα-GFP for control; AAV-CaMKIIα-cre-mCherry for test group; titer, 4.8 × 10^12^ GC/ml for both). Surgery was performed on adult floxed-*Kcna2* mice (8 weeks). After anesthesia with isofluorane, mice were secured in a stereotaxic frame (Neurostar, Tubingen, Germany). Holes were drilled bilaterally in the skull at the injection sites (4 sites). Stereotaxic coordinates used for intrahippocampal injections were as follows (from bregma); for a dorsal hippocampus, anterior–posterior: -2.1 mm, mediolateral: ±2.2 mm, dorsoventral: 2.25 mm; for a ventral hippocampus, anterior–posterior -2.85 mm, mediolateral ±2.8 mm, dorsoventral 3.3 mm. A 33 gauge needle attached to a 50 μl Hamilton syringe (#80908), mounted to the stereotaxic frame, and under control of a microinjection syringe pump UMP3T-1 controller (WPI, Sarasota, FL, USA) was used to inject 0.4 μl of AAV at each site. Injections occurred at a rate of 0.04 μl/min, after which the needle was left in place for an additional 2 min. After injections were completed, the skin was sutured and the animals were allowed to recover for 1 h on a heating pad before returning to the home cage. Mice remained in the homecage for an additional 4 weeks before the start of behavioral testing and post-hoc *ex vivo* experiments.

### Preparation of slices for electrophysiological recording

Acute transverse hippocampal slices were obtained from Sprague-Dawley rats (P15-P22; P, postnatal days) or aforementioned mice (P15-P24) of either sex. Animals were anesthetized by inhalation of 5 % isoflurane. After decapitation, brain was quickly removed and chilled in ice-cold cutting solution contained 75 mM sucrose, 87 mM NaCl, 25 mM NaHCO_3_, KCl 2.5 mM, NaH_2_PO_4_ 1.25 mM, D-glucose 25 mM, MgCl_2_ 7 mM, and CaCl_2_ 0.5 mM (equilibrated with carbogen mixture of 95% O_2_ and 5% CO_2_). After mounting, 300 μm-thick transverse slices were prepared in the cutting solution using a vibratome (Leica VT1200, Nussloch, Germany), and incubated at 34 °C for 30 min in the same solution, and thereafter stored at room temperature (22 °C). For experiments, slices were transferred to a submersion recording chamber superfused with standard recording solution contained 124 mM NaCl, 26 mM NaHCO_3_, 3.2 mM KCl, 1.25 mM NaH_2_PO_4_, 10 mM D-glucose, 2.5 mM CaCl_2_, and 1.3 mM MgCl_2_.

### Electrophysiological recordings

Whole-cell voltage- or current-clamp recordings from CA3-PCs were performed at near-physiological temperature (34 ± 1 °C) in standard recording solution while the recording chamber was perfused with the recording solution at 1 ∼ 1.5 ml/min. Patch pipettes were pulled from borosilicate glass tubing (Outer diameter: 1.5 mm, Wall thickness: 0.225 mm) with a horizontal pipette puller (P-97, Sutter Instruments, Novato, CA, USA) and filled with the intracellular solution contained 130 mM K-gluconate, 7 mM KCl, 1 mM MgCl2, 2 mM Mg-ATP, 0.3 mM Na-GTP, 10 mM HEPES, 0.1 mM EGTA (pH = 7.20 with KOH, 295 mOsm with sucrose), and pipette resistance was 3 ∼ 4 MΩ. Recordings were preferentially obtained from the hippocampal CA3b. After formation of whole-cell configuration on the somata of CA3-PCs, recordings were performed from only cells that had stable resting membrane potential within -76 and -58 mV (cells that had more positive resting membrane potential or unstable resting membrane potential was discarded). Under this condition, input conductance (G_in_) was measured from subthreshold voltage responses to -30 pA and +10 pA current steps with 0.5 sec (Hyun et al., 2015). G_in_ was monitored every 10 s before and after the activities delivered to CA3-PCs described below. The first spike latency (AP latency) was measured from voltage responses of CA3-PCs to a ramp current injection (250 pA/s for 1 s) (Hyun et al., 2013). All recordings were made using a MultiClamp 700B amplifier controlled by Clampex 10.2 through Digidata 1400A data acquisition system (Molecular Devices, Sunnydale, CA, USA)

### Synaptic stimulation for MF-, and PP-CA3 synapses

Various stimulations were delivered to the synapses of CA3-PCs were delivered to evaluate the influence to the intrinsic excitability of CA3-PCs. Afferent MFs were stimulated with a recording solution-filled glass monopolar electrode with resistance of 1 ∼ 2 MΩ placed in striatum (st.) lucidum (SL; stimulus intensity with 2 ∼ 20 V) using minimal stimulation techniques (Hyun et al., 2015). Afferent PP and fibres were stimulated with a concentric bipolar electrode (CBAPB125; FHC Inc., Bowdoin, ME, USA). Brief stimulation pulses (100 μs) was generated by a digital stimulator (DS8000, WPI; Sarasota, FL, USA) and delivered to stimulation electrode through an isolation unit (DLS100 stimulus isolator, WPI). A stimulation electrode was positioned at st. lacunosum-moluculare (SLM) on the border of the subiculum and CA1 (for PP stimulation), and identified the type of synaptic inputs by the rise time of EPSCs and sensitivity to group II mGluR agonist, (2S,2’R,3’R)-2-(2’,3’-Dicarboxycyclopropyl)glycine (DCG-IV, 2 μM). The attenuation of PP-EPSCs and MF EPSCs were 72.8%, 72.6 % (data not shown), consistent with previous reports (Tsukamoto et al., 2003).

### In situ hybridizaion

Fluorescence in situ hybridization (FISH) for RNA transcripts was performed using RNAscope probes [Advanced Cell Diagnostics (ACD), Hayward, CA, USA] for target genes following the manufacturer’s instructions. A harvested brain was quickly frozen on liquid nitrogen. Frozen sections of 15 ∼ 20 μm thick were serially and coronally cut through the habenula formation. Sections were thaw-mounted onto Superfrost Plus Microscope Slides (Fisher Scientific #12-550-15). The sections were fixed in 4% PFA for 10 min, dehydrated in increasing concentrations of ethanol for 5 min, and finally air-dried. Tissues were then pretreated for protease digestion for 10 min at room temperature. For hybridization, prepared slides were incubated in HybEZ hybridization oven (ACD) for 30 min at 40°C. Unbound hybridization probes were removed by washing the sections three times with 1x wash buffer at room temperature for 2 min. For signal amplification, slides were incubated with amplification solutions (Amplifier *#*-FL): #1 for 30 minutes at 40°C, #2 for 15 min at 40°C, #3 for 30 minutes at 40°C, #4 for 15 min at 40°C. After the completion of each session, slides were washed twice with 1x wash buffer for 2 min at room temperature. Nuclei were counterstained with 4′,6-diamidino-2-phenylindole (DAPI, ACD). For detection of *Homer1a (H1a)*, *Arc*, and *Kcna2* mRNA in Figure 4 and Figure 5, RNAscope Probes Mm-Homer1-tvS (Cat#, 433941), Mm-Arc-C3 (Cat#, 316911-C3), Mm-Kcna2-C2 (Cat#, 462811-C2) were used, respectively. The slides were viewed, analyzed, and photographed using TCS SP8 Dichroic/CS (Leica, Germany) or Olympus Fluoview FV1200 (Japan) confocal microscope equipped with 488 nm and 633 nm diode lasers. For *H1a* and *Arc*, the settings for photomultiplier and laser power were optimized for detection of intra-nuclear foci and for minimizing weaker cytoplasmic signals (Almira et al., 2002). The number of *Kcna2* mRNA dots and intra-nuclear foci of *H1a* and *Arc* were counted on 60 x magnified photographs (N.A. = 1.2).

### Image acquisition and analysis for catFISH

Hippocampal slices obtained from *WT*, *Kcna1+/−*, and *Kcna2+/−* mice were analysed with RNAscope assay (Advanced Cell Diagnostics, Hayward, CA, USA) using target probes for Arc (Cat No.: 540901) and Homer1a (Cat No.: 433261) for catFISH described in Figure 4. Confocal z-stacks composed of 1-μm-thick optical sections were collected in regions CA3 from 3 slides/animal and stored for offline analysis. Stacks for CA3 (one stack per slide) were collected with a 20× objective to collect comparable numbers of cells from each subjects. Non-neuronal cells, identified as small cells (c.a. 5 μm in diameter) with intensely bright and uniformly stained nuclei, were excluded from the analysis. Only large, diffusely stained nuclei present in the sections were regarded as neuronal cells, and included in our analysis. Neuronal nuclei from the CA3 region were classified as negative (containing no transcription foci), H1a (+) (containing only H1a transcription foci), Arc (+) (containing only Arc transcription foci), or Arc/H1a (+) (containing transcription foci for both Arc and H1a) by an experimenter blind to the relationship between the image stacks and the behavioural conditions they represented. The number of nuclei corresponded to each classification and total number of neuronal nuclei were counted.

### Evaluation of CA3 region-specific Kv1.2 hetero-knockout mice

Hippocampal slices obtained from CA3-Kcna2+/− and floxed-Kcna2 mice were analysed with RNAscope assay (Advanced Cell Diagnostics, Hayward, CA, USA) using RNAscope^®^ Probe-Mm-Kcna2-C2 (Cat No.: 462811-C2). Confocal z-stacks composed of 1-μm-thick optical sections were collected in regions CA3 from 3 slides/animal and stored for offline analysis. The number of mRNA dot in CA3, CA1, and dentate gyrus (DG) region was counted by an experimenter blind to the relationship between the image stacks and the behavioural conditions they represented, based on an assumption that each RNA dot deriving from a single mRNA molecule (Wang et al., 2012).

### Contextual fear discrimination test

We tested 39 male mice between 14 and 20 weeks of age (20 WT mice, 10 kcna1+/– and 9 kcna2+/– mice) for fear conditioning in a pair of similar contexts using a protocol adapted from (McHugh et al., 2007). This test assesses an animal’s ability to discriminate between two similar contexts, A and B, through repeated experience of a foot shock that is associated with Context A but not with B. Context A (conditioning context) is a chamber (18 cm wide × 18 cm long × 30 cm high; H10-11M-TC; Coulbourn Instruments 5583, PA 18052, USA) consisting of a metal grid floor, aluminium side walls, and a clear Plexiglass front door and back wall. The context A chamber was lit indirectly with a 12 W light bulb. The features of Context B (safe context) were the same as Context A, except for a unique odor (1% acetic acid), dimmer light (50% of A), and a slanted floor by 15° angle. Each chamber was cleaned with 70% ethanol before the animals were placed. On the first 3 days of the experiment, the mice were placed in Context A, where they were allowed a 3-min exploration, received a single foot shock (1 mA, for 2 s) and were returned to their home cage 60 s after the shock. Freezing levels were measured during the 3 min before the shock delivery. On day 4, mice of each genotype were divided into two groups; one group of genotype visited Context A and the other Context B. On day 5, we had each mouse visit the context opposite to the one visited on day 4, and freezing levels were measured again. On day 4-5, neither group received a shock in Context A and B. The mice were subsequently trained to discriminate these two contexts by visiting the two contexts daily for 9 days (discrimination task day 6 to 14), always receiving a footshock 3 min after being placed in Context A but not B (Figure 3A). The freezing ratio (measured during the first 3 min) in each context was used to calculate discrimination ratios for each animal in both groups over the 9 days of training. We defined freezing behavior as behavioral immobility except for movement necessary for respiration and assessed the freezing behavior of each mouse for the duration of 5 min (day 4-5) or 3 min (day 6-14) by observing its video image for 2 s bouts every 10 s and counting the number of 2 s bouts during which the mouse displayed freezing behavior (referred to as a freezing score). Freezing ratio was calculated as the freezing score divided by the total number of observation bouts (18 or 30). Discrimination ratios were calculated according to F_A_ / (F_A_ + F_B_), where F_A_ and F_B_ are freezing scores in Contexts A and B, respectively. All experiments and analyses were performed blind to the mice genotype.

### One-trial contextual fear conditioning

We tested five WT mice and five *Kcna2*+/− mice between 14 and 16 weeks of age for fear conditioning in a pair of very distinct contexts using the experimental schedule of 3 days (acclimation, conditioning, assessment) according to (Cravens et al., 2006). The aforementioned Context A was used as the conditioning context. For the distinct context (Context C), we put a white acrylic blind end cylinder (15 cm in diameter, 18 cm in height, and 0.5 cm in thickness) vertically on the metal grid floor of the conditioning chamber, and covered the bottom inside the cylinder with cage bedding, on which mice were placed. The chamber and cylinder were cleaned using 70% ethanol between runs. On day 1, we first placed mice in Context A and then placed them in Context C an hour later. Mice were allowed to freely explore in both contexts for 5 min. On day 2, we had mice revisit Context A and receive a single foot shock (1 mA, for 2 s) 3 min later, and returned them to their home cage 60 s after the shock. On day 3, mice were separated into two groups; mice of each group were placed in Context A or C for 3 min without a foot shock, during which the freezing score was measured. All experiments were conducted and analysed by scientists blind to the genotypes of the mice.

### Pre-exposure-mediated contextual fear conditioning (PECFC)

This task requires mice to retrieve contextual memory from a very brief exposure to a previously experienced context (Fanselow, 1990; Nakashiba et al. 2008). Male WT mice (n = 18) and male Kcna2+/− (n = 18) mice between 15 to 25 weeks of age were tested for PECFC. The pre-exposure context was the same as the aforementioned ‘Context A’, whereas non-exposure context was the same as the aforementioned ‘Context C’. The chamber was cleaned with 70% ethanol between runs. On day 1, each group of genotypes was divided, and one subgroup were allowed to freely explore Context A for 10 min, and the other group were allowed to Context C for same time (pre-exposure). On the second day, each group were separated into conditioned and unconditioned groups. The conditioned subgroup mice were placed into Context A for 10 s, received a single foot shock (1 mA, for 2 s) and were immediately (30 s after the shock) returned to their home cages. Mice of the unconditioned subgroup were just brought back to the home cages 42 s after being placed in the Context A without a foot shock. On day 3, 24 h after experiencing Context A, mice of each group were placed in Context A and a freezing score was assessed for 3 min.

### Elevated plus maze (EPM)

We assessed basal anxiety level in WT mice (n = 5) and kcna2 HT mice (n = 10) between 13 and 16 weeks of age using the elevated plus maze. The apparatus consisted of two open (30×5 cm) and two closed (30 × 5 × 15 cm) arms facing each other with an open roof. The entire maze was elevated at a height of 40 cm and each animal was placed individually on the central platform (5 × 5 cm), facing an open arm, and was allowed to explore the apparatus for 5 min. Anxiety was measured by the time spent in open arms. Data were collected using a video camera fixed to the ceiling of the room and connected to a video tracking equipment and a recorder using EthoVision software (Noldus Information Technology, Wageningen, Netherlands).

### Statistical Analysis

Statistical data are expressed as mean ± standard error of the mean (SEM), and the number of cells/animals measured (denoted as n; details were described in results). Statistical data were evaluated for normality and variance equality with Kolmogorov-Smirnov test and Levene’s test, respectively. For data that satisfy normality and equality of variances, statistical evaluations were performed with student’s t-test or ANOVA. For data that inappropriate for parametric tests, non-parametric tests were performed for evaluation. The number of cells and statistical tests for determining statistical significance are stated in the text using following abbreviations: n.s., no statistical significance; ∗, P < 0.05; ∗∗, P < 0.01; ∗∗∗, P < 0.005. Statistical analyses were performed using PASW Statistics 18 (SPSS Inc, 2009, Chicago, IL, USA).

## Acknowledgements

This study was supported by grants from National Research Foundation of Korea (Grant No. 2020R1A2C2006438) and Seoul National University Hospital.

---

**Figure 1-figure supplement 1.**
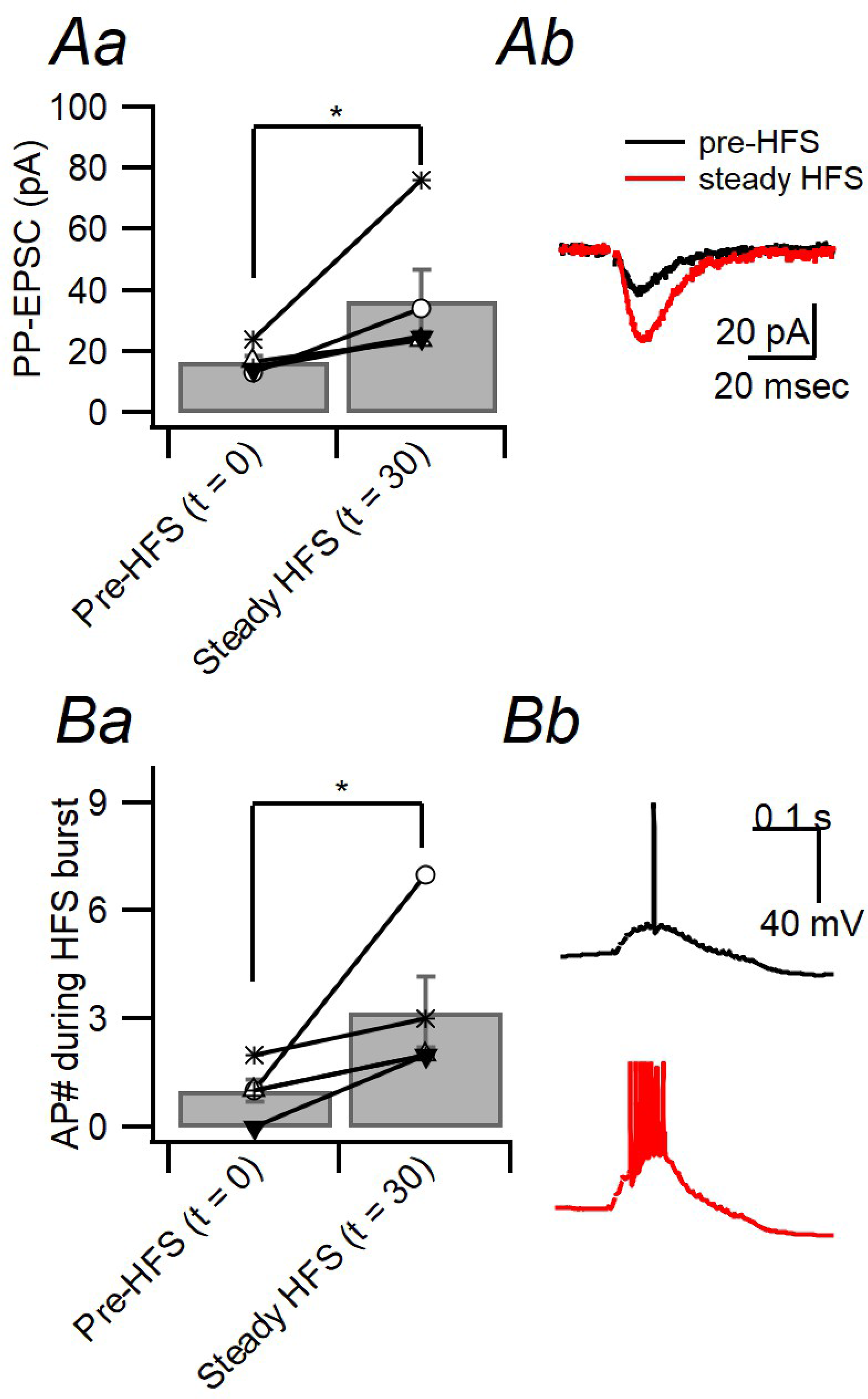
PP-LTP enhances the number of spikes in CA3-PCs elicited by a burst stimulation of PP facilitates the generation of AP spikes of CA3-PCs. **Aa**. potentiation of PP-EPSCs before and after PP-HFS (the same protocol as shown in Figure 1Ab (n = 5, Z = −2.06, p = 0.039, Wilcoxon signed rank test). **Ab**. Exemplar traces for PP-EPSCs before (black; t = −5 min) and after (red; t = 30 min) PP-HFS. **Ba**. In the same PP-CA3 synapses (denoted by the same symbols as in A), the number of APs by a single burst stimulation (20 pulses at 100 Hz) of PP-CA3 synapses was significantly increased after induction of PP-LTP (n = 5, Z = −2.06, p = 0.039). **Bb**. Exemplar traces for temporally summated PP-EPSPs and APs evoked by a single burst stimulation before (upper, black) and after (lower, red) the induction of PP-LTP.

**Figure 2-figure supplement 1.**
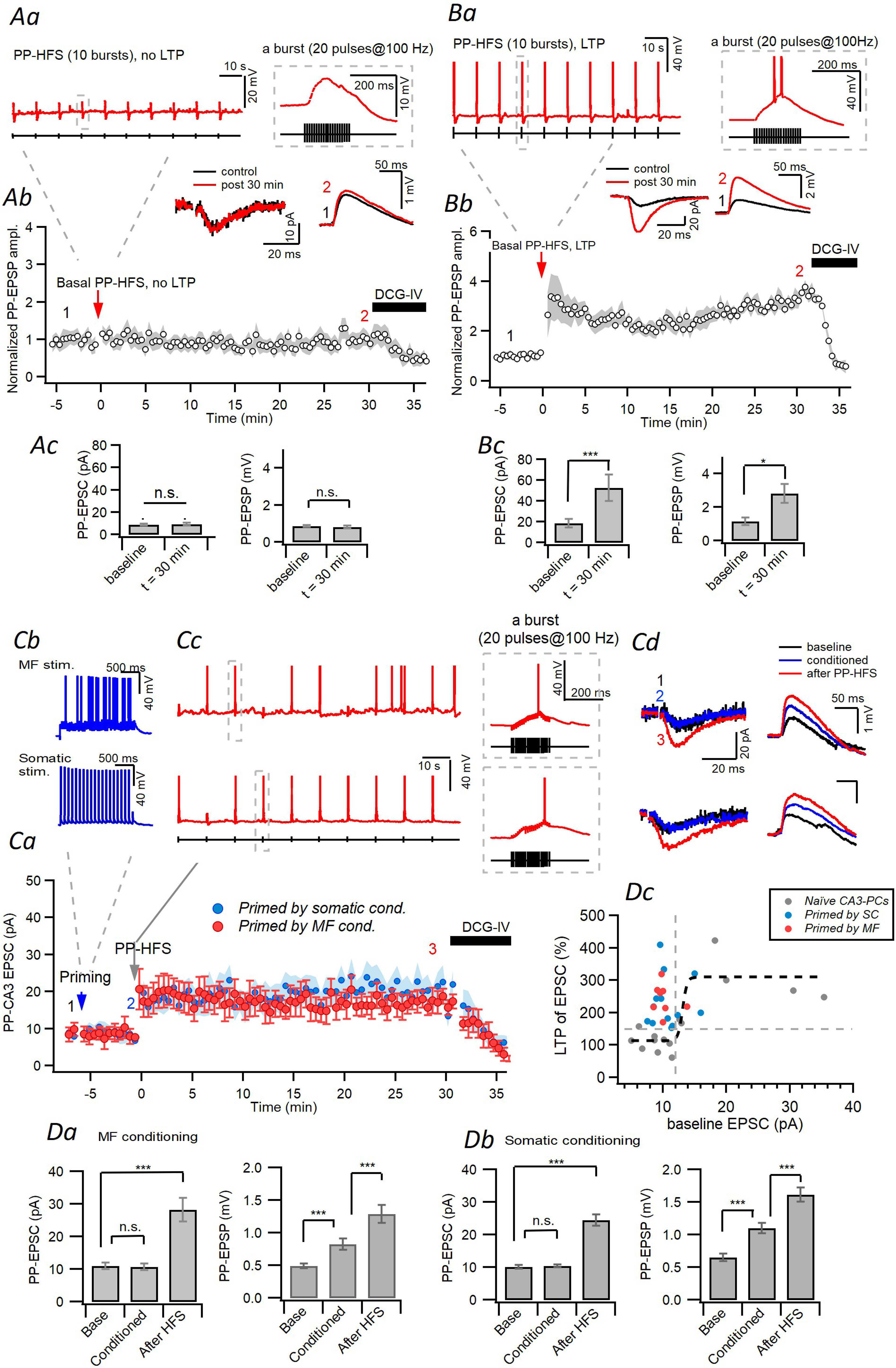
Both 10 Hz somatic stimulation and 20 Hz MF stimulation lower the threshold for PP-LTP induction in the rat CA3-PCs. **Aa.** Weak PP-HFS protocol (*black*) and postsynaptic voltage responses (*red*). The pulse protocol for PP-HFS is the same as in Figure 1. *Right inset* corresponds to the gray-dashed box on the left. **Ab.** Weak PP-HFS (baseline PP-EPSC < 12 pA) induced no potentiation of PP-EPSPs. *Insets*, Representative traces for PP-EPSCs (*left*) and PP-EPSPs (*right*) before (*black*) and 30 min after HFS (*red*) at PP-CA3 synapses. EPSCs were measured at the start and the end of EPSP monitoring. **Ac.** Mean amplitude of EPSCs (*left*) and EPSPs (*right*) before and 30 min after PP-HFS. Under this weak stimulation intensity conditions, the baseline amplitudes for EPSPs and EPSCs of non-failure events were 0.92 ± 0.08 mV and 10.24 ± 1.64 pA, respectively. No significant change of EPSCs and EPSPs was induced by weak PP-HFS in the naïve CA3-PCs (EPSP, t = 0.264, p = 0.799, n = 9; EPSCs, t = −0.620, p = 0.555, n = 8; independent t-test, measured at 30 min). **Ba.** Strong PP-HFS protocol. The pulse protocol is the same as the weak version (*Aa*) but higher stimulation intensity. Note that a burst of APs was elicited by strong HFS in CA3-PCs. **Bb.** Normalized EPSPs monitored at PP-CA3 synapses. *Insets*, Representative PP-EPSC and PP-EPSP traces before (*black*) and 30 min after HFS (*red*). Both EPSCs and EPSPs became larger after strong PP-HFS. **Bc**. Mean amplitudes of EPSCs (*left*) and EPSPs (*right*) before (Pre) and 30 min after HFS. Both EPSCs and EPSPs were potentiated by strong PP-HFS (EPSC, n= 5, t = −7.312, p = 0.002; EPSP, n = 5, t = −4.133, p = 0.014; independent t-test). **Ca.** Monitoring of PP-EPSCs before and after 20 Hz MF stimulation (*red*) or 10 Hz somatic APs (*blue*) for 2 s, and subsequent weak PP-HFS. Note that EPSCs are not altered by conditioning, but potentiated by weak PP-HFS. **Cb.** CA3-PCs were primed by induced by consists of MF stimulation at 20 Hz for 2 s (*upper*) or repetitive somatic firing at 10 Hz for 2 s (*lower*). Each current injection is amplitude of 1200 pA, duration of 2.2 ms. **Cc.** Diagram (black) and response (red) of weak HFS after the MF stimulation (*upper;* number of elicited APs for 2 s, 27.5 ± 3.44) or somatic firing (*lower*). HFS consisted of 10 bursts delivered per 10 s, and each bursts consisted of 20 stimuli at 100 Hz (gray-dashed line box of *right*, corresponded to gray-dashed line box of *left*). This HFS protocol was applied at -60 mV, adjusted in current-clamp mode. The stimulation intensity of this HFS is sub-threshold, which is insufficient to induce LTP without conditioning (Refer to the Figure 1). **Cd**. Representative EPSCs (left) and EPSPs (right) traces before and after MF (*upper*) or somatic (*lower*) conditioning. *black*, baseline; *blue*, after conditioning; *red*, after PP-HFS. **Da-Db.** Summary for mean amplitudes of EPSCs (left) and EPSPs (right). Note that no change in EPSCs after MF conditioning (*Da*, t = −0.725, p = 0.492, n = 8) and somatic conditioning (*Db*, n = 13, t = −0.012, p = 0.991; independent t-test), while EPSPs were potentiated by MF conditioning (*Da*, n = 8, t = −5.262, p = 0.001) and somatic conditioning (*Db*, n = 13, t = −9.409, p < 0.001, independent t-test). Weak PP-HFS subsequent to MF conditioning increased EPSCs (*Da*, n = 8, t = −9.027, p < 0.001), and EPSPs (*Da*, n = 8, t = −6.380, p < 0.001, independent t-test). Similarly, HFS after somatic conditioning also increased EPSCs (*Db*, n = 13, t = −9.027, p < 0.001) and EPSPs (*Db*, n = 13, t = −9.613, p < 0.001, independent t-test). **Dc.** LTP of PP-EPSCs as a function of PP stimulation intensity quantified as the baseline EPSC amplitude. PP-LTP in CA3-PCs conditioned by MF (red) or somatic firing (blue) is compared to that in naïve CA3-PCs (gray). Note that the induction threshold for PP-LTP is lowered by conditioning.

**Figure 3-figure supplement 1.**
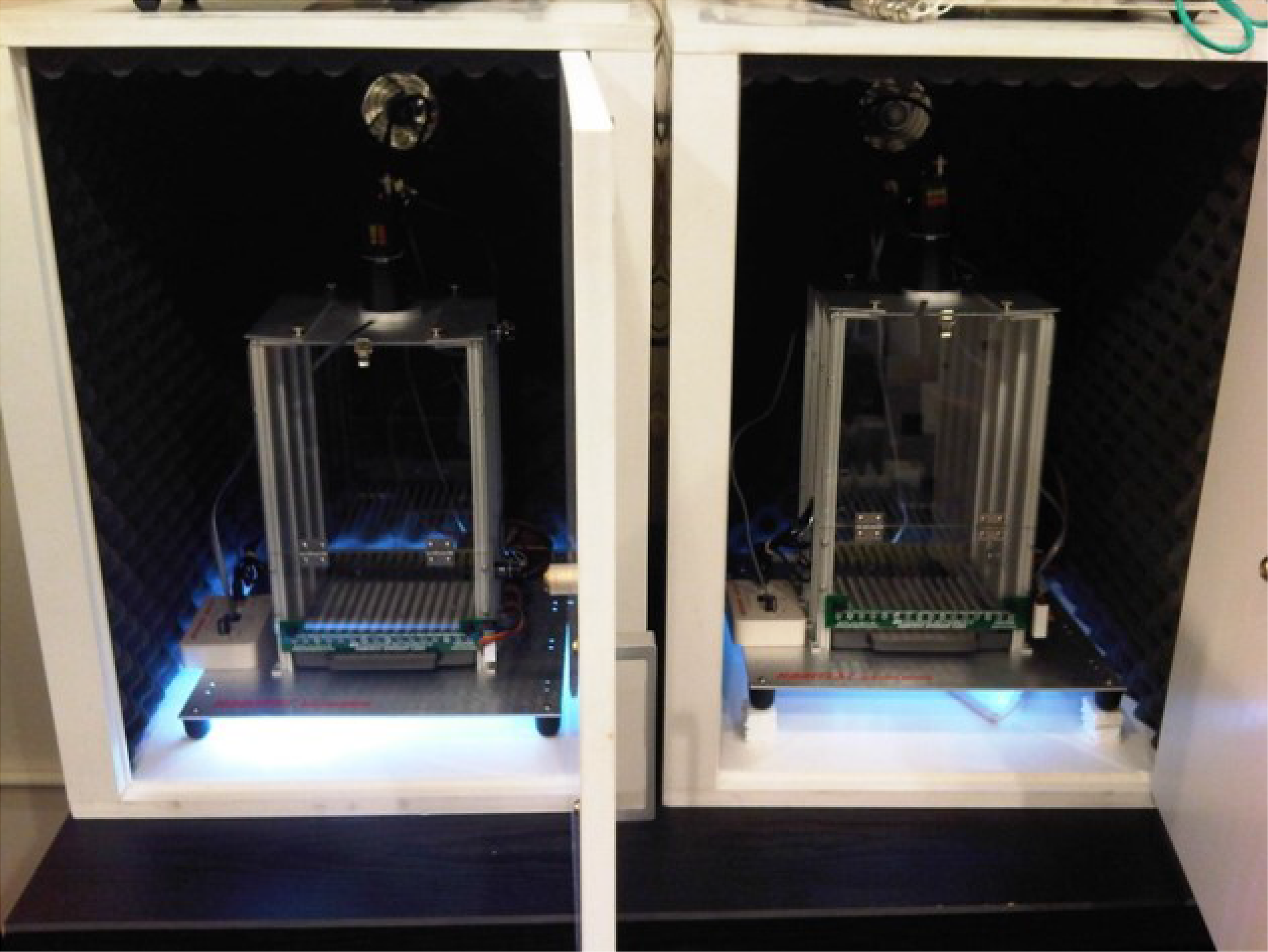
Photographs of contexts A and B. The context A (left) is a chamber consisting of a metal grid floor, aluminium side walls, and a clear Plexiglass front door and back wall. The context A chamber was lit indirectly with a 12 W light bulb. The context B (right) had the same sound noise level, metal grid floor, sidewalls and roof as in context A, but differed from context A in having a unique odor (1% acetic acid), dimmer light (50%) and a slanted grid floor by 15° angle.

**Figure 4-figure supplement 1.**
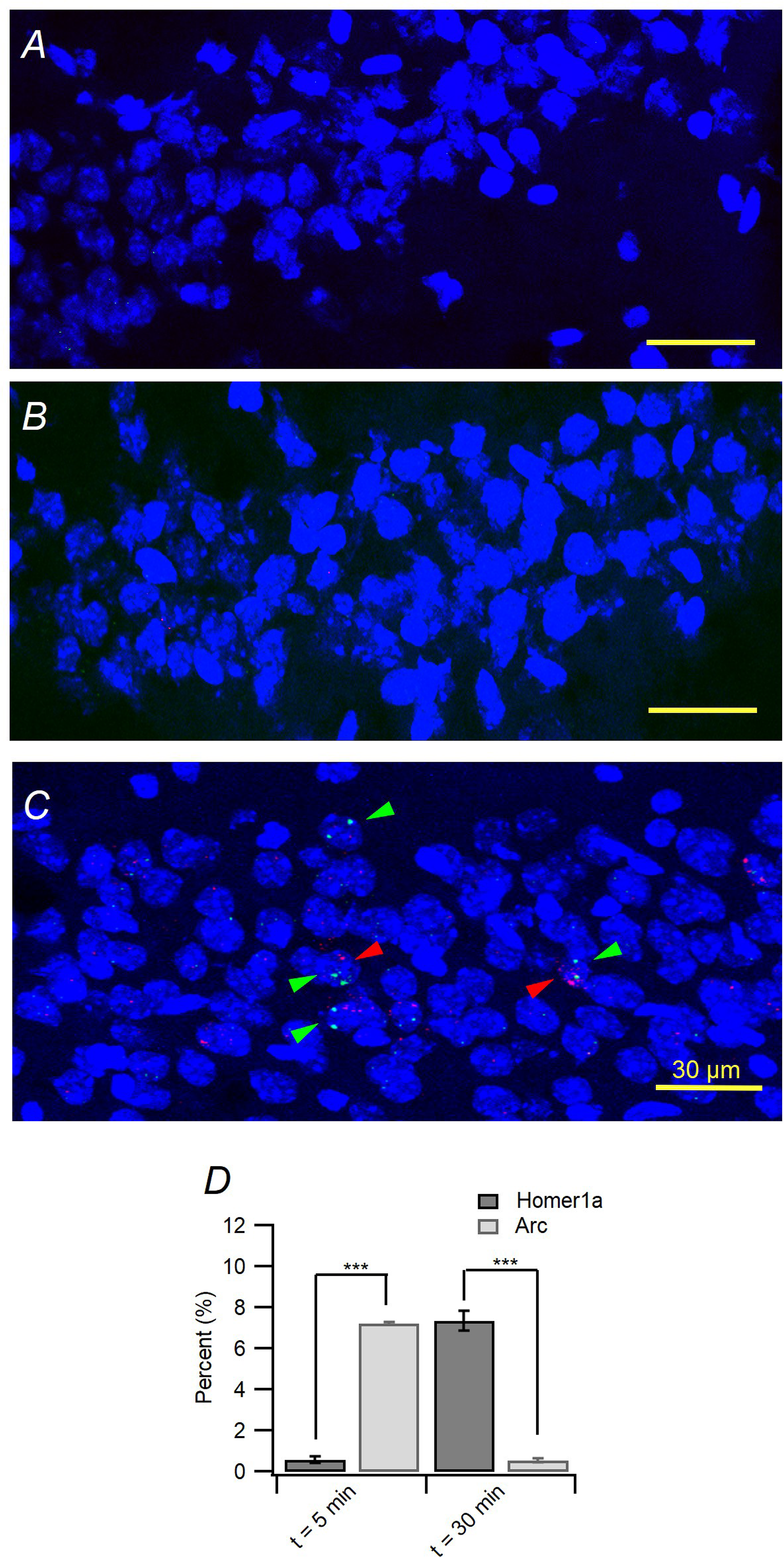
Test for non-specific expression of H1a and Arc INF. **A-B**, To evaluate non-specific expression of H1a and Arc transcripts, catFISH for H1a (*A*) and Arc (*B*) mRNA was performed at 5 min and at 30 min, respectively, after the mice being placed in a novel context (context A) with a footshock. Scale bars, 30 μm. **C.** catFISH image reproduced from Figure 4Aa for comparison. Green and red arrowheads indicate H1a and Arc INF. **D**. Mean fractions of H1a and Arc-positive nuclei, when each was detected at 5 or 30 min after the mice were exposed to a novel context. Note that most nuclei were negative for H1a and Arc INF, when H1a and Arc mRNA were stained at 5 and 30 min after the mice visited a novel context (H1a INF: 7.34 ± 0.49% at 30 min, 0.54 ± 0.01% for 5 min, Mann-Whitney U = 0.00, p = 0.004, n =4; Arc INF: 7.21 ± 0.37% at 5 min, 0.60 ± 0.02% for 30 min, Mann-Whitney U = 0.00, p = 0.004, n =4).

**Figure 4-figure supplement 2.**
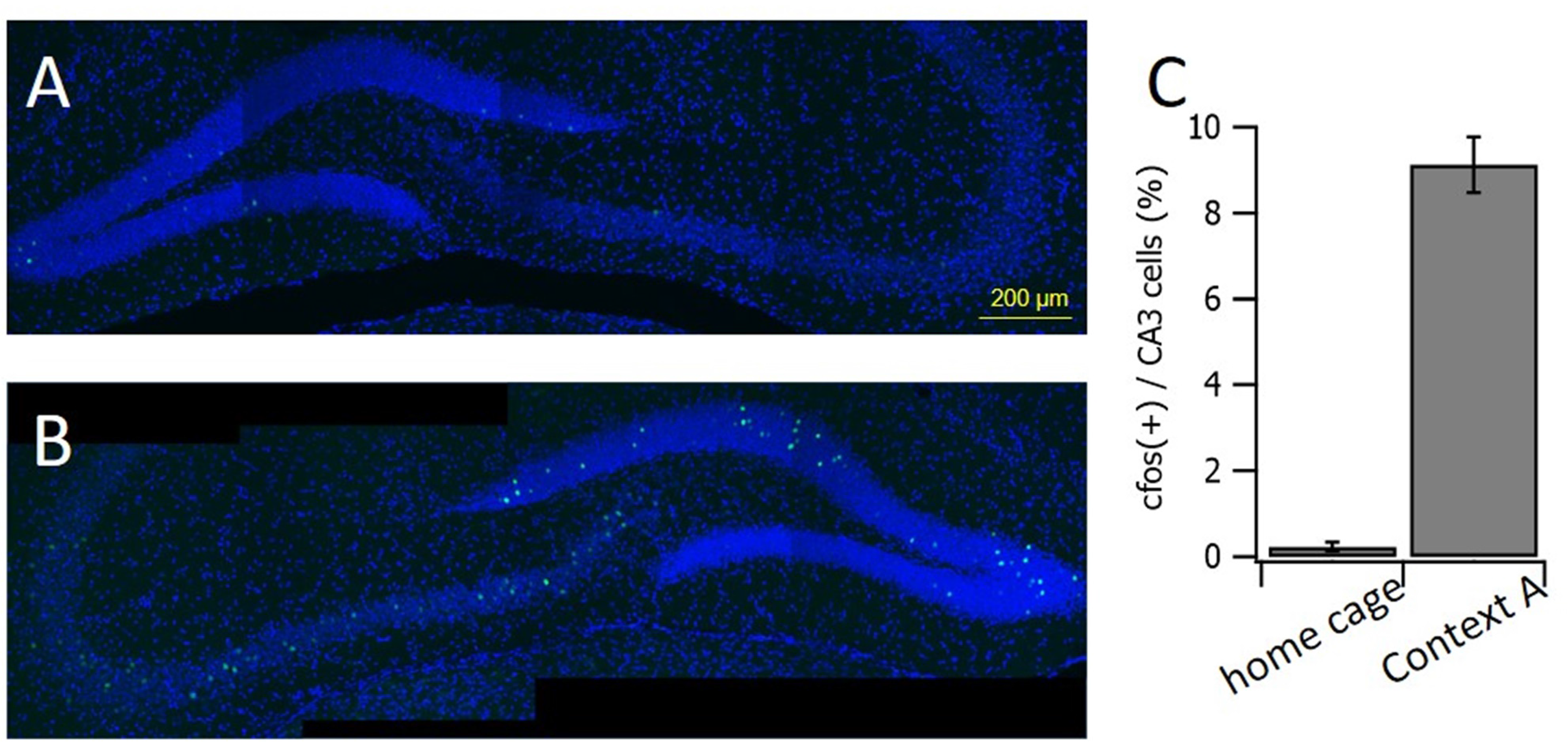
**A-B**, Representative fluorescence images of c-fos in the dorsal hippocampi from WT mice, which have been handled for a week. On day 8, the mice were killed immediately from their home cage (*A*) or 30 min after being placed in a novel context (context A) with a footshock (*B*), and then their hippocampi underwent immunostaining for c-fos (green) with DAPI counterstaining of nuclei (blue). Antibody targeting c-fos and DAPI were purchased from Cell Signaling (Cat# 2250, USA) and Sigma-Aldrich (St Louis, MO, USA), respectively. **C**, Proportions of c-fos(+) nuclei in the CA3 area of mice from homecage (n = 4) and those from context A (n = 5 slices, two mice for each).

**Figure 6-Figure Supplement 1.**
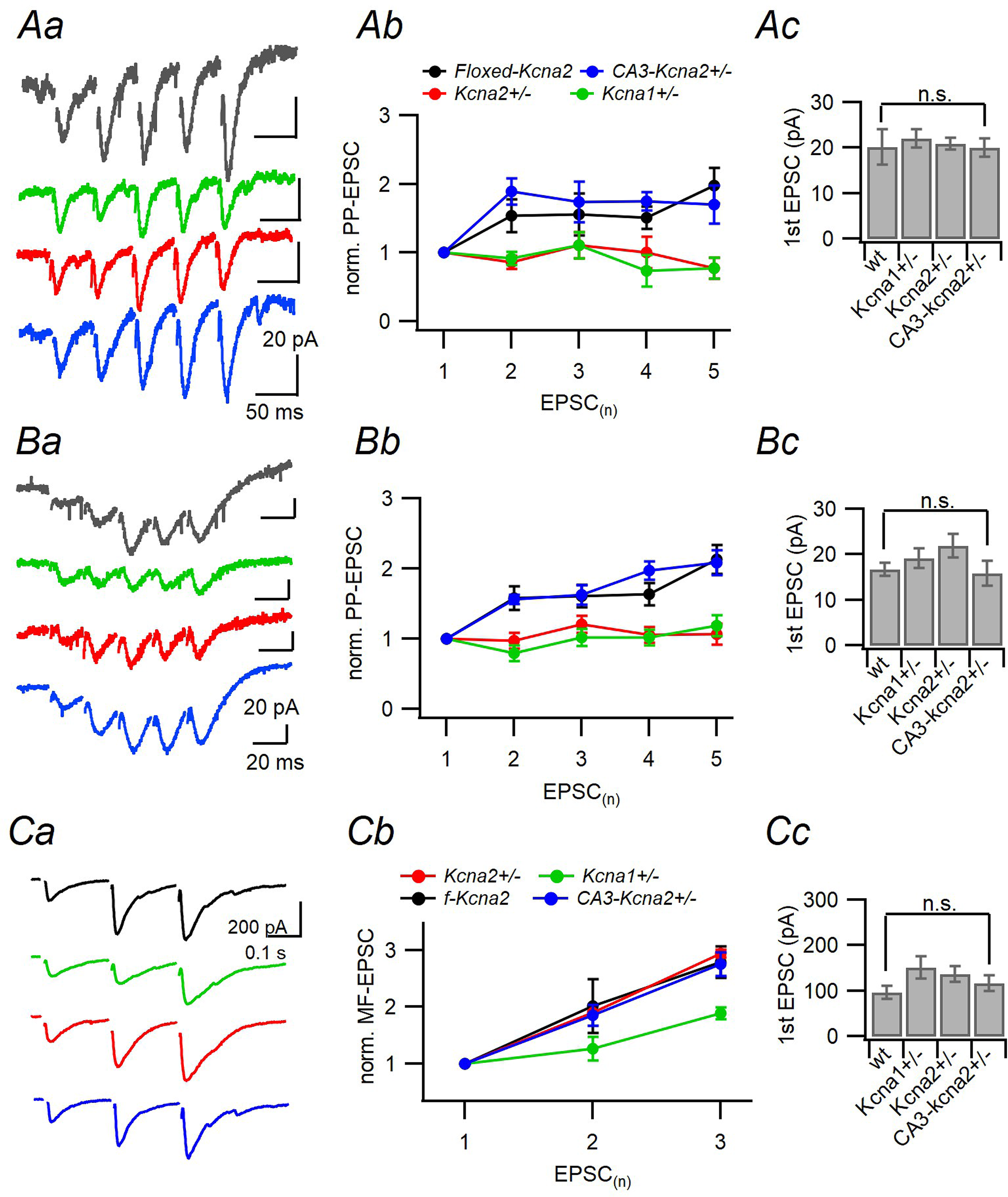
Short-term plasticity of PP and MF-CA3 synapses. **Aa, Ba and Ca**, Representative traces for EPSCs evoked by 20 (*Aa*) or 50 Hz (*Ba*) PP stimulation and 50 Hz MF stimulation (*Ca*) in CA3-PCs of *f-Kcna2* (black), CA3-*Kcna2*+/− (blue), *Kcna1*+/− (green), and *Kcna2*+/− (red) mice. **Ab, Bb and Cb**, Relative amplitudes of EPSC trains normalized to the 1^st^ EPSC evoked by 20 Hz (*Ab*) or 50 Hz (*Bb*) PP stimulation or 50 Hz MF stimulation (*Cb*). **Ac, Bc and Cc**, Mean amplitude of the 1st EPSCs in different genotypes. No significant difference in the 1^st^ EPSC amplitudes between genotypes, because the afferent fiber stimulation intensity was adjusted such that the 1^st^ EPSC amplitude falls within a narrow range. wt, *f-Kcna2*; n.s., statistically not significant.

